# Fitness costs of noise in biochemical reaction networks and the evolutionary limits of cellular robustness

**DOI:** 10.1101/068510

**Authors:** J. David Van Dyken

## Abstract

Gene expression is inherently noisy, but little is known about whether noise affects cell function or, if so, how and by how much. Here I present a theoretical framework to quantify the fitness costs of gene expression noise and identify the evolutionary and synthetic targets of noise control. I find that gene expression noise reduces fitness by slowing the average rate of nutrient uptake and protein synthesis. This is a direct consequence of the hyperbolic (Michaelis-Menten) kinetics of most biological reactions, which I show cause “hyperbolic filtering”, a process that diminishes both the average rate and noise propagation of stochastic reactions. Interestingly, I find that transcriptional noise directly slows growth by slowing the average translation rate. Perhaps surprisingly, this is the largest fitness cost of transcriptional noise since translation strongly filters mRNA noise, making protein noise largely independent of transcriptional noise, consistent with empirical data. Translation, not transcription, then, is the primary target of protein noise control. Paradoxically, selection for protein-noise control favors increased ribosome-mRNA binding affinity, even though this increases translational bursting. However, I find that the efficacy of selection to suppress noise decays faster than linearly with increasing cell size. This predicts a stark, cell-size-mediated taxonomic divide in selection pressures for noise control: small unicellular species, including most prokaryotes, face fairly strong selection to suppress gene expression noise, whereas larger unicells, including most eukaryotes, experience extremely weak selection. I suggest that this taxonomic discrepancy in selection efficacy contributed to the evolution of greater gene-regulatory complexity in eukaryotes.

**ARTICLE SUMMARY:** Gene expression is a probabilistic process, resulting in random variation in mRNA and protein abundance among cells called “noise”. Understanding how noise affects cell function is a major problem in biology. Here I present theory demonstrating that gene expression noise slows the average rate of cell division. Furthermore, by modeling stochastic gene expression with non-linearity, I identify novel mechanisms of cellular robustness. However, I find that the cost of noise, and therefore the strength of selection favoring robustness, decays faster than linearly with increasing cell size. This may help explain the vast differences in gene-regulatory complexity between prokaryotes and eukaryotes.

## INTRODUCTION

The probabilistic nature of chemical reactions, along with the small volumes of living cells, create substantial molecular randomness called “noise” (Berg 1978; McAdams and Arkin 1997). Gene expression noise, for example, is measured as variation in mRNA or protein abundance within a single cell over time (Taniguchi et al. 2010), or, equivalently, among cells in an isogenic cell population (Elowitz et al. 2002). Because the abundance of most mRNAs and proteins are on the order of 1 to 1000 copies per cell (Taniguchi et al. 2010), a random birth or death of a single copy can instantaneously change the cellular concentration by 0.1-100%. These random fluctuations are amplified by bursting, the synthesis of multiple mRNAs or proteins in brief pulses (Raser and O’Shea 2004; Raser and O’Shea 2005; Raj et al. 2006; Pedraza and Paulsson 2008; Thattai and van Oudenaarden 2001). Using fluorescent probes and fluorescently tagged proteins, researchers have quantified the noise statistics of mRNA and/or proteins in *E. coli,* budding yeast, and mammalian cell lines, demonstrating the ubiquity of gene expression noise and its considerable variability across the genome and among taxa (Elowitz et al. 2002; Blake et al. 2003; Golding et al. 2005; Raj et al. 2006; Zenklusen, Larson, and Singer 2008; Taniguchi et al. 2010; Newman et al. 2006; Balázsi, van Oudenaarden, and Collins 2011).

A central question in cell biology and evolution is whether or not cellular noise has a functional consequence. Answering this question is necessary for establishing rational design criteria for synthetic genetic circuits and for providing evolutionary explanations for empirical patterns of noise among genes and taxa. For example, why does *E. coli* have an average mRNA Fano factor (variance/mean) of *F* = 1.6 (Taniguchi et al. 2010), which is just above the theoretical minimum of *F* = 1, whereas mammalian cells have 40-100 times stronger mRNA noise (Raj et al. 2006; Balázsi, van Oudenaarden, and Collins 2011; Dar et al. 2012)? While the mechanistic causes of this discrepancy are becoming clearer (Suter et al. 2011), the evolutionary forces permitting such a discrepancy remain murky.

Mounting empirical evidence suggests that natural selection has acted to limit gene expression noise in many cases (Metzger et al. 2015; Newman et al. 2006; Alemu et al. 2014; Batada and Hurst 2007; Fraser et al. 2004; Lehner 2008), implying that noise is often costly to cells. Yet identifying and quantifying this cost remains a major challenge. Perhaps the most widely invoked cost of cellular noise is imprecision, the lack of fine control over cellular processes. Evolutionarily, this cost of noise is typically modeled as random phenotypic deviations from the adaptive optimum, which is opposed by stabilizing selection (Wagner et al. 1997; Draghi and Whitlock 2015). Several molecular mechanisms linking imprecision to suboptimal fitness deviations have been proposed.

The dosage-control hypothesis proposes that noise disrupts control over the stoichiometric balance of interacting proteins (Fraser et al. 2004; Lehner 2008). Consistent with this hypothesis, proteins that form complexes have lower average noise than other proteins (Fraser et al. 2004; Lehner 2008; Bar-Even et al. 2006), but models currently do not predict the magnitude of this effect so that quantitative correspondence between predictions and observations remain unknown. More commonly, researchers focus on how noise disrupts the fine control over cell states regulated by bistable (ON/OFF) switches (Balázsi, van Oudenaarden, and Collins 2011; Csete and Doyle 2002). Noise in the input parameter can cause the cell to flip stochastically from the desired to undesired state, or to respond slowly to environmental cues, creating a maladaptive mismatch between the cell state and the selecting environment. However, the least noisy genes, those presumably experiencing the strongest selection to suppress noise, are constitutively expressed and involved in core cell functions, not control over bistability (Fraser et al. 2004; Lehner 2008; Bar-Even et al. 2006). This cost is also difficult to quantify in any general way because its effects are highly context dependent, to the extent that in some contexts the cost of imprecision can change signs, becoming evolutionarily beneficial (Acar, Mettetal, and van Oudenaarden 2008; Rao, Wolf, and Arkin 2002; Kussell and Leibler 2005; Eldar and Elowitz 2010; Kaern et al. 2005; Thattai and Van Oudenaarden 2004).

As an alternative, here I demonstrate a general, readily quantifiable and consistent cost of cellular noise. I show that noise in substrate abundance slows the average rate of product formation in Michaelis-Menten-type (i.e., non-cooperative bimolecular) chemical reactions, which constitute most reactions in the cell including those most closely tied to fitness: nutrient uptake and protein synthesis. This result is a straightforward consequence of the non-linearity of bimolecular reactions. While it is well known that noise alters the mean of nonlinear stochastic systems, this effect is typically ignored, giving rise to the widespread use of linear approximations (e.g., the “linear noise approximation” (LNA) (van Kampen 2007)), particularly in studies of gene expression noise (Thattai and van Oudenaarden 2001; Shahrezaei and Swain 2008; Paulsson 2004). The LNA is well-justified when applied to investigating steady-state concentrations in noisy chemical networks, since the difference between linear and non-linear predictions at steady-state are on the order of a single molecule. However, fitness in living cells is determined by rates—not concentrations--integrated over an enormous number of reactions over the lifetime of a cell and its genetic lineage. For example, an *E. coli* cell must carry out a minimum of 2.5x10^6^ translation reactions to produce a single daughter cell. A minor slowing of reaction rates within cells, if heritable, may pose a non-negligible selective cost. From this perspective, the effects of noise on fitness cannot be ignored, and indeed, I find that they can be substantial in small cells.

## RESULTS

The goal of the paper is to quantify the strength of selection favoring the attenuation of gene expression noise, and to identify the targets of noise attenuation. The paper proceeds as follows. First, I show that noise slows the average rate of biochemical reactions that have non-linear, hyperbolic kinetics. Then, I extend this result to networks of coupled non-linear reactions in order to determine how noise effects total flux through biochemical networks, as well as how the network architecture itself affects the propagation of noise. This enables us to quantify the cost of noise, measured as the loss in average rate of end-product formation, and to quantify the noise statistics of the system. I then apply the model to gene expression, focusing on the consequences of non-linear, hyperbolic translation kinetics on the propagation of mRNA noise to the protein level, and to the average rate of biomass synthesis. Finally, I investigate how the volume of reaction compartments and whole-cells influences the cost of noise, concluding that the cost of noise, and therefore the selection pressures favoring low gene expression noise, decays rapidly with increasing cell size, leading to a testable prediction about levels of gene expression noise across unicellular species.

### Noise-induced slowdown of Michaelis-Menten reactions

The purpose of this section is to begin building the theory from the simplest component of a network: a single isolated reaction. We will see that input noise slows the reaction rate of a single, isolated reaction obeying Michaelis-Menten type kinetics. In the following section, the analytical results from this section are put on a more rigorous theoretical footing and then extended to networks of coupled reactions in open systems.

The cornerstone of biochemistry is the Michaelis-Menten (MM) reaction, which describes the enzyme catalyzed conversion of substrate, *S*, into product, *P*. The rate of product formation is described by the hyperbolic MM equation (Michaelis and Menten 1913; Briggs and Haldane 1925; Ingalls 2013):

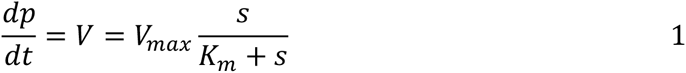

*s, e* and *p* are reactant, enzyme and product concentration, respectively (following convention lowercase denotes concentration, which unless otherwise noted will take units of molecules per cell), *V_max_ = k_cat_e* is the asymptotic reaction rate (*k_cat_* is the number of substrate molecules converted to product per enzyme per unit time), and *K_m_*, the Michaelis or half-saturation constant, is the substrate concentration at *V* = *V_max_*/2. This equation is derived from a system of elementary reactions by applying the law of mass action and the quasi-steady state approximation (QSSA) (Briggs and Haldane 1925; Ingalls 2013) (Sup. Mat.). The QSSA is a separation of timescales approximation that eliminates fast variables by setting them to their steady-state. The law of mass action is valid in cases where the timescale of molecular diffusion is fast relative to the reaction rate, so that the system behaves as if it were well-mixed. The rate of enzymatic reactions in cells is far below the diffusion limit (Bar-Even et al 2006), so that mass action is a reasonable approximation in most cases. For stochastic reactions, wherein *s, e*, and *p* experience random number fluctuations, Eqn. 1 takes on a probabilistic interpretation as the “reaction propensity” (van Kampen 2007) giving the transition probability for *n_P_* ➜ *n_P_* + 1 and *n_S_* ➜ *n_S_* – 1, where *n* denotes particle number. In general, the QSSA is valid for stochastic systems under the same conditions for which it is valid in deterministic systems (Rao and Arkin 2003; Sanft, Gillespie, and Petzold 2011), notwithstanding finite-volume corrections (Grima 2009b; Grima 2009a; Grima 2010) (see below).

Because Eqn. 1 is hyperbolic in *s*, Jensen’s inequality proves that, for random *s*, 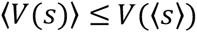 (angle brackets denote ensemble average). That is, the average reaction rate is less than or equal to a noise-free (i.e., deterministic) reaction with *s* = 〉*s*〈 (Figs. 1, S1). This is seen by expanding Eqn. 1 to second order in Taylor series about 〉*s*〈 and taking the expectation (see SI),

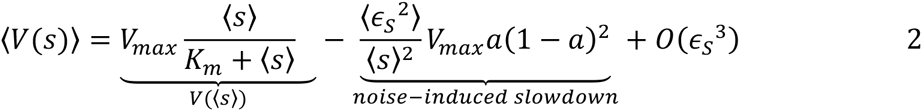

*a* = *K_m_*/(*K_m_* + 〉<u>s</u>〈) is the “kinetic order” of the MM reaction evaluated at the average substrate concentration, and 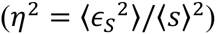 the squared coefficient of variation, which is often simply called “the noise”. The leading order correction in Eqn. 2 takes the sign of the second derivative of Eqn. 1, which is everywhere negative for hyperbolic transition rates, thus demonstrating the noise-induced slowdown of MM type reactions.

The kinetic order parameter, *a*, is a key parameter that appears repeatedly below. *a* is identical to the steady-state fraction of unbound enzyme. In metabolic control analysis (MCA), *a* is equivalent to the “elasticity coefficient” (Kacser and Burns 1973). In general, *a* measures the nonlinearity of the reaction rate at a given mean substrate concentration. First and zeroth order reactions (*a* = 1 and 0, respectively) are both linear, and as a consequence there is no noise-induced slowdown, consistent with Jensen’s Inequality. The rate of a first order reaction is linearly dependent on the substrate concentration, whereas the rate of a zeroth order reaction is independent of substrate concentration. In a MM type reaction, the reaction is in its first order regime when substrate is scarce, and in the zeroth order regime when substrate is abundant (the flat part of the rate law curve in Fig. 1).

**Figure 1:**
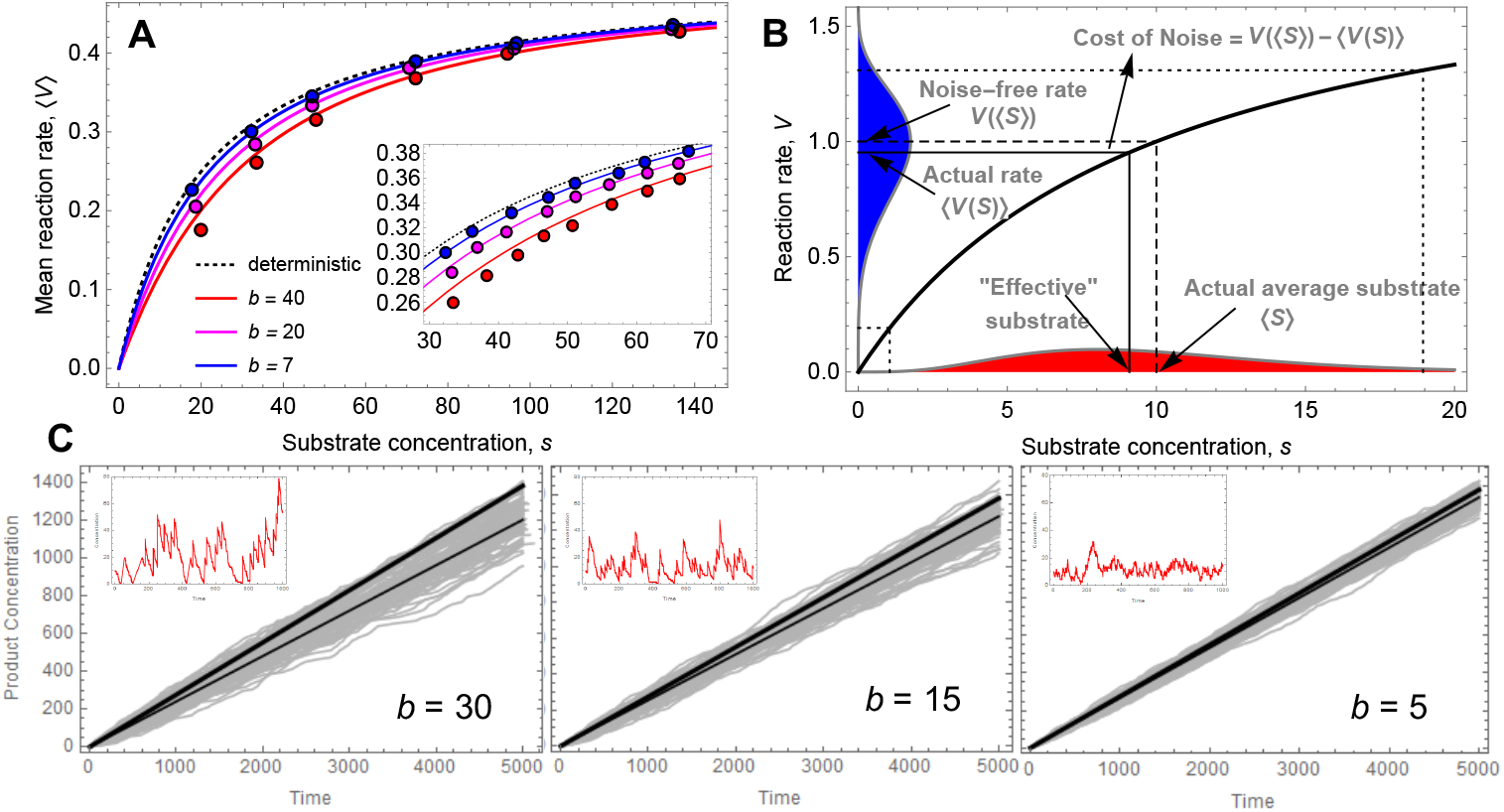
Noise-induced slowdown of hyperbolic reactions. Stochastic simulations employing Gillespie’s (Gillespie 1977) exact stochastic simulation algorithm (SSA) of the microscopic system of elementary reactions for the Michaelis-Menten reaction (see SI). Points represent the rate of product formation over time steps 100-800 (providing a burn-in more than 10 times the half life of *P* or *S*) ensemble averaged over 1000 realizations of the stochastic process. Results fit well with the QSSA model (Eqn. 3, solid lines in **a, b** and thin black line in **c**). The substrate burst input, *b*, is varied to alter the substrate noise level. Substrate input rate, *k_in_* is divided by *b* so that mean substrate is not affected by bursting, and is varied to generate different substrate concentrations. All other parameters are constant (*e* = 1, *k_ON_* =1, *k_OFF_* = 19.5, *k_cat_* = 0.5, δ_S_ = 0.1, δ_P_ = 0.1). **A**) The rate law curve demonstrating that, for a given average substrate concentration, substrate noise slows the reaction, but only if the reaction is not in its first or zeroth order regime (low or high substrate, respectively). Note that the accuracy of the stochastic QSSA requires that *s*, *K_m_* <<*F*, which fails for the red points in **a** and **b** (where *K_m_* = 20 and *F* = 20.5). Inset is a zoomed in view. **B**) A geometric illustration of noise-induced slowdown, showing that variance (noise) in a state variable reduces the average of a hyperbolic function (i.e., Jensen’s inequality). **C**) 100 realizations of the stochastic process (gray) compared to the deterministic (thick black line) and stochastic (thin black line) predictions. Red traces in inset are example outputs from a single realization of the substrate dynamics for a short time window.

A more rigorous and general expansion method is used in the following section (“Noisy Non-linear Reaction Networks in Finite Volumes”). For now, there are two main points: 1) reaction kinetics depend not only on average molecule concentrations, but on the mean and variance in molecule numbers, and 2) for a given average substrate concentration, substrate noise reduces the rate of biochemical reactions. The difference between the average stochastic rate and deterministic rate grows monotonically with increasing noise, and equality occurs only with zero noise or zero curvature of *V*(*s*). Mechanistically, this can be understood as follows: positive fluctuations in substrate oversaturate available catalysts leaving many unbound *S* molecules that are not actively creating product, causing inefficiency. As a consequence, the same amount of substrate would produce a faster net reaction rate if spread out evenly over the same time interval (i.e., the same mean with zero noise).

If the substrate statistics are known, then the probability density of *V* with random *s* can be solved explicitly for a number of common probability distributions. mRNA and protein counts are typically gamma distributed within a cell over time or among cells in a clonal population (Taniguchi et al. 2010). The gamma distribution has shape parameter α = *CV*^−2^ and rate parameter *β* = *F*, where 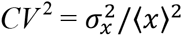 and 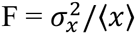 are the squared coefficient of variation (the noise) and the Fano factor (often called “the noise strength”), respectively (Taniguchi et al. 2010). *F* and *CV*^2^ are common measures of noise intensity, with convenient mechanistic interpretations in gene expression: *F* is closely related to the burst size (e.g., number of proteins synthesized per mRNA lifetime) and *CV*^−2^ to the burst frequency (e.g., the number of mRNA’s synthesized per protein lifetime). *F* is independent of cell volume, whereas *CV*^2^ decreases with cell volume (they are intensive and extensive variables, respectively). For gamma distributed *s*, the probability density function of the Michaelis-Menten reaction rate (for *V* 〉 *V_max_)* is exactly,

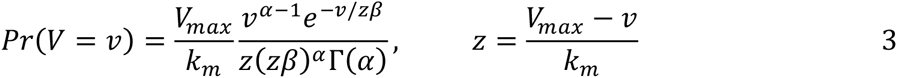

Eqn. 3 is verified by Monte Carlo simulations implemented with Gillespie’s exact stochastic simulation algorithm (SSA) (Gillespie 1977) (Fig. 1). Eqn. 3 fits data from stochastic simulations very well provided that *F, K_m_* ≪ *s*. Equation 2 gives a very good approximation to 3 under this same restriction (Fig. S2), which provides an important benchmark for the accuracy of the more rigorous expansion approximations used below.

Figure 1 shows that the effect of noise on MM-type reactions is to increase the observed Michaelis constant. Therefore, a stochastic formulation of Eqn. 1 that accounts for noisy substrate while retaining the classical form is obtained simply by substituting into Eqn. 1 an “effective”, stochastic Michaelis constant, 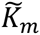, where 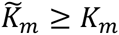 with equality occurring only with zero noise, zero curvature of the rate law, and macroscopic reaction volume (S6).

### Noisy Non-linear Reaction Networks in Finite Volumes

In the previous section, we showed that substrate noise slows the rate of product formation in a single hyperbolic reaction in a closed system. To apply this result, the statistics of the substrate species are required. We now take a more rigorous approach and generalize the noise-induced slowdown phenomenon to reaction networks in open systems. Further, the model allows one to predict the noise statistics of the system solely with knowledge of the network structure and reaction kinetics. Of particular interest is how network architectures may create feedbacks that buffer the system flux against noise-induced slowdown. The following theory builds on the work of van Kampen (van Kampen 2007) and Elf and Erhenberg (Elf and Ehrenberg 2003) (see also Grima (Grima 2010)).

The time evolution of the joint probability density of all *N* nodes in a chemical reaction network with *R* edges is given by the following “system-size expansion” (van Kampen 2007) of the multidimensional chemical master equation (derived in the SI):

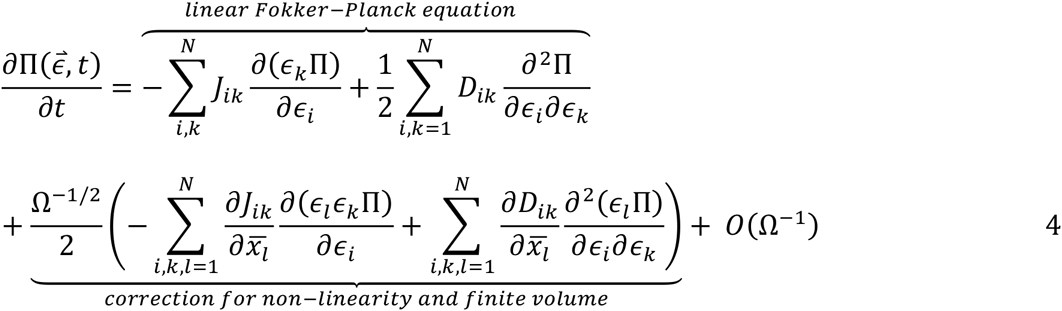

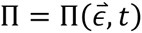 is the joint probability density of all reacting species, Ω is the reaction volume, *ϵ_i_* is the random perturbation of the *i*^th^ species from 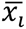, its macroscopic steady-state (overbars will denote the macroscopic steady-state value of a variable throughout), *S_ij_* is the stoichiometric coefficient of species *i* in the *j*^th^ reaction, and

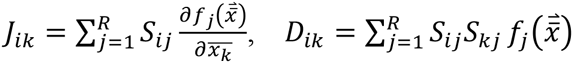

are entries of the Jacobian and diffusion matrices, respectively, of the continuous deterministic system of equations. Equation 4 is a linearization of the chemical master equation obtained by applying the ansatz that concentrations fluctuate about their macroscopic steady-state value, with fluctuation size scaling inversely with the square root of the reaction volume, Ω (van Kampen 2007). This inverse square-root dependence of noise on volume is a formal statement of the Law of Large Numbers, and is formally equivalent to the relationship between sample size and the standard deviation of a sampling distribution. One thus substitutes, 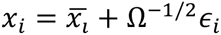, for each species *i* in the deterministic system of equations, and then expands each transition rate, *V_j_(x)*, in powers of Ω^−1/2^ about 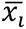, truncating the series at the desired order (van Kampen 2007). The terms of order *O*(Ω^0^) in (4) give the “linear noise approximation” (LNA) (van Kampen 2007), which is equivalent to the standard linear Fokker-Planck equation, while the terms in Ω^−1/2^ account for noise-induced deviations caused by non-linear rate laws in finite volumes, which we show cannot be ignored in living cells.

To apply 4, one begins by writing a macroscopic description of a reaction network in QSSA reduced form. Here we analyze a generalized MM scheme (S17) in an open system obeying the following macroscopic system of ODEs (Sup. Mat.):

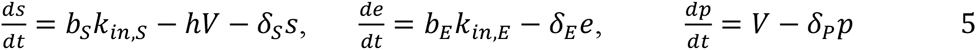

*δ_i_* and *k_in,i_* are the decay and input rates, respectively, of *i*, and *V* = *k_cat_es*/(*K_m_*+*s*). Substrate (*S*) and enzyme (*E*) are fed into the system via a burst process with burst size *b_s_* and *b_E_*, respectively 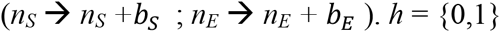 is an indicator variable that toggles between reactions where *S* is consumed (*h* = 1), as in most metabolic reactions, and reactions where *S* is not consumed (*h* = 0), as with template-mediated reactions such as translation where *E* denotes ribosomes and *S* mRNA, which is not consumed in translation.

From System 5 and Eqn 4, we obtain the following steady-state average concentrations of each species in a generalized, open MM reaction network (Sup. Mat.):

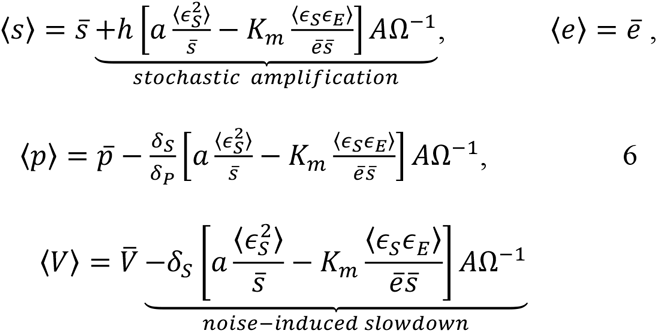

Where 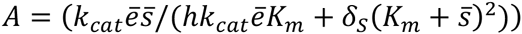 and 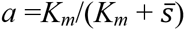 is the MM reaction order. Note that in the general case, 〉*s*〈 is increased and 〉*p*〈 is decreased by noise compared to the macroscopic system. The non-linear mesoscopic correction terms vanish when *s*➔0 (first order kinetics), *s*➔Infinity (zeroth order kinetics) or when *Ω*➔ Infinity (macroscopic volume). Two of the second moments (Sup. Mat.) for the general MM case can be written compactly:

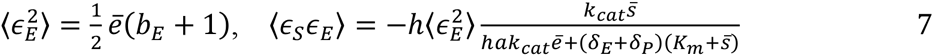

While the present approach builds on that derived by Grima (Grima 2010) the results and conclusions differ: Grima (Grima 2010) found that noise had no effect on product concentration in MM reactions. There are two reasons for this discrepancy. First, Grima assumed no dilution of the substrate (*δ_s_* = 0). As can be seen from equation 6, this causes the finite volume correction term for 〉*p*〈 to vanish. Thus, Grima’s results are in agreement with the present results for this special case, and all of the results in his papers are correct. However, following (Grima 2010) exactly for *δ_s_* > 0 also gives zero stochastic deviation of the product concentration (see SI), which is incorrect. Here is why: when the system-size expansion of the chemical master equation is applied to a system of elementary reactions, as in (Grima 2010) and most other methods, the rate of product formation, *V*, is proportional to the intermediate complex concentration, *c*. Because *c* is unimolecular, the method reads this rate as linear (its second derivative is zero), and so the mean cannot be affected by noise (as proven by Jensen’s inequality). However, in MM reactions *V* is decidedly non-linear. Indeed, it is hyperbolic. Thus, the elementary reaction conceals an essential non-linearity of the system, and will give incorrect results in the general case. To rectify this, one must apply Eqn. 4 to a QSSA reduced system of biochemical equations as shown in the SI. Note that the use of elementary or QSSA reactions is not up to choice: the elementary reaction scheme will give the wrong answer. Note also that applying the method of (Grima 2010) to a QSSA reduced system also gives incorrect results, for reasons discussed in the SI.

### Noise propagation in gene expression with hyperbolic filtering

Because gene expression noise is costly, there will ostensibly be selection pressure for noise control. But how can a cell suppress gene expression noise? That is, what are the evolutionary targets of noise-control? To answer these questions, we must model the process of gene expression, form transcription through translation, to identify how noise is generated and filtered.

We deviate from previous models of gene expression noise by assuming that translation obeys non-linear reaction kinetics. Previous models of gene expression noise have considered translation as a first-order process (Thattai and van Oudenaarden 2001; Shahrezaei and Swain 2008; Paulsson 2004; Bar-Even et al. 2006; Pedraza and Paulsson 2008; McAdams and Arkin 1997). However, translation in living cells may actually be closer to zeroth-order than first-order, with a reaction order of *a* = 0.1-0.2 in log-phase growth in *E. coli* and budding yeast, based on the fraction of unoccupied ribosomes (Arava et al. 2003; Zenklusen, Larson, and Singer 2008; Ingolia 2014). We now investigate the effects of hyperbolic translation kinetics on gene expression noise.

Following previous models of gene expression noise (Thattai and van Oudenaarden 2001; Shahrezaei and Swain 2008; Paulsson 2004; Bar-Even et al. 2006; Pedraza and Paulsson 2008; McAdams and Arkin 1997), we assume for now that translation is initiation limited (see “Cell fitness” section below for generalization), such that it relies only on the concentrations of ribosomes, *r*, and mRNA, *m*. If mRNA is transcribed in bursts of size *b_t_*, then the steady-state variance and Fano factor of mRNA are (Sup. Mat.),

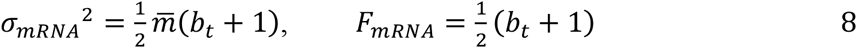

which is a special case of (Elf and Ehrenberg 2003) and which is in excellent agreement with simulations (Figs 2, 3).

**Figure 2:**
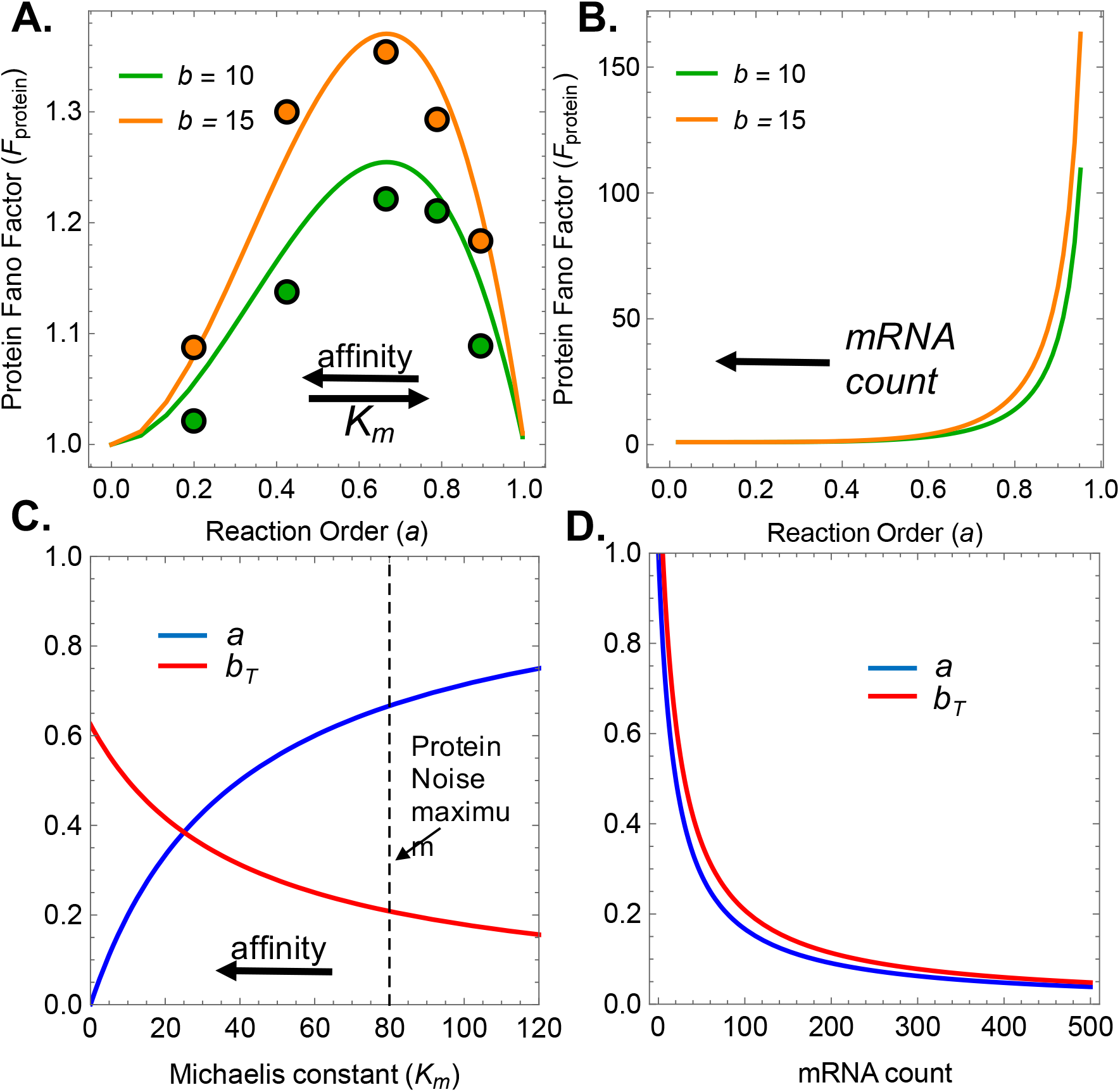
Protein noise with hyperbolic filtering. Translation propagates mRNA noise to the protein level, but hyperbolic reaction kinetics (0<α<l) can substantially ameliorate this noise and cause highly non-linear relationships between noise and underlying parameters. Points represent ensemble variance of protein number from the last of 110 time steps over 10,000 realizations of the SSA (see SI), lines represent theoretical prediction (from Eqns 9 and S21). **A, C**) Increasing ribosomal binding affinity (i.e., reducing K_m_) has two opposing effects on protein noise: it reduces reaction order but increases translational bursting. Which effect dominates depends on the population’s starting point relative to the protein noise peak at a* = 2/3. Most species have translational *a* < 2/3, suggesting that increasing ribosomal binding affinity will actually reduce protein noise, even though it increases bursting. **B, D**) Increasing mRNA abundance, holding mRNA noise constant, monotonically decreases protein noise for all parameter values, as both the reaction order and translational burst parameters decrease with increasing mRNA (Sup. Mat. for theoretical derivation of this result). Parameters: e = 5, k_ON_ =1, k_OFF_ =19.5 (varied in **A**), k_cat_ = 0.5, δ_S_ = 0.1, δ_P_ = 0.1, k_in_ = 4/b (varied in **B**). K_m_ was tuned by varying k_OFF_, s was tuned by varying k_in_, while mRNA noise was held constant with a burst input of either b =10 or 15.

**Figure 3:**
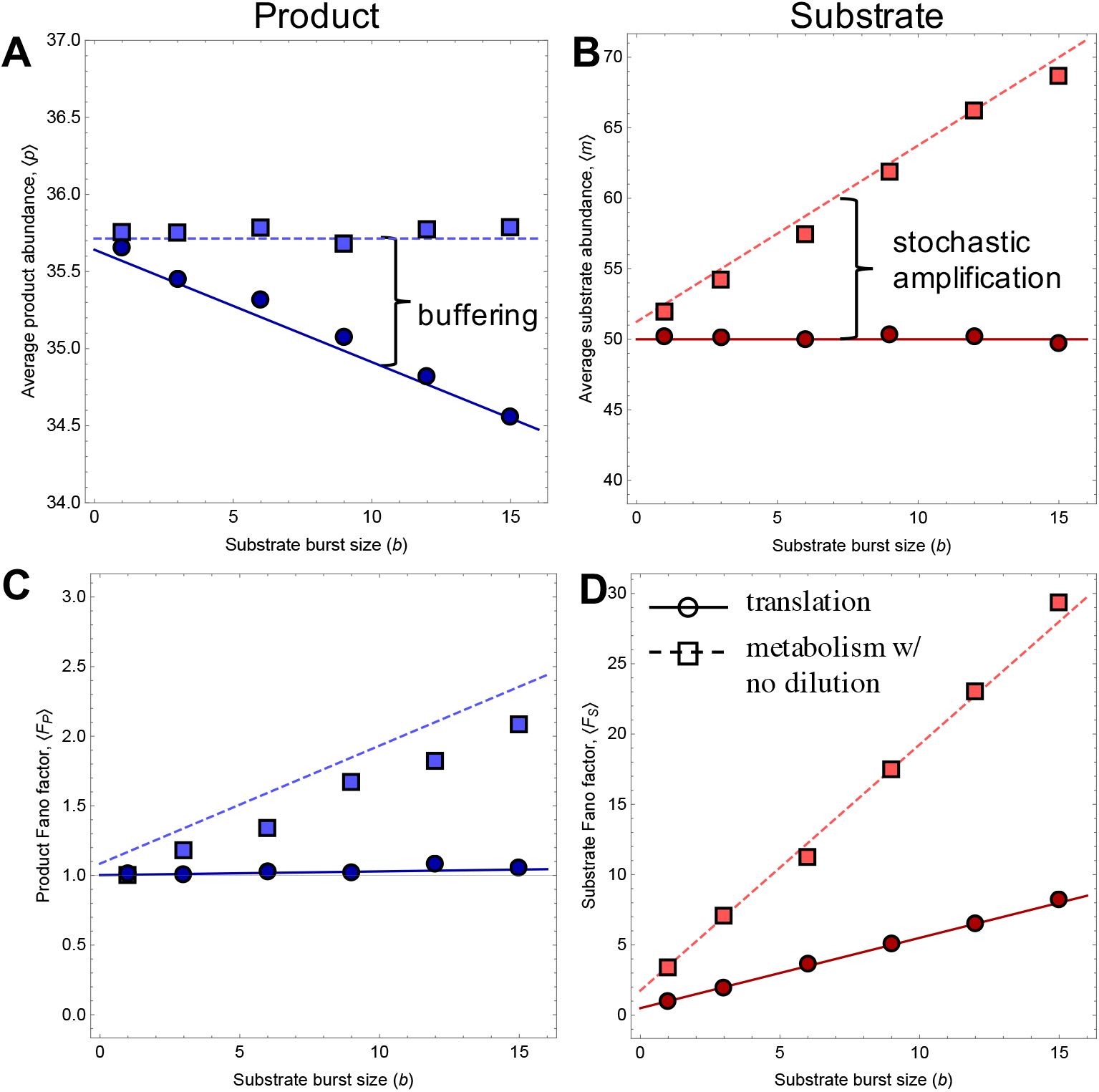
Effect of network architecture on the cost of noise. Theoretical predictions (Eqns. 6–8, S20, S21) for metabolic reactions (dashed lines; *h* = 1, δ_s_ = 0) and translation (solid lines; *h* =0, δ_s_ = 0.1) compared to ensemble average (blue) and variance (red) of 1,000-5,000 SSA realizations (points). **A**) Noise-induced slowdown causes an accumulation of substrate (“stochastic amplification”) which completely buffers reaction flux for metabolic reactions when substrate dilution/degradation is zero (δ_S_ = 0) (**B**). Because mRNA is not consumed in translation (*h* = 0), its concentration is independent of reaction rate so does not experience stochastic amplification. Thus, translation is not buffered by network feedback. **C,D**) The effect of network architecture on substrate (C) and product (D) noise. e = 1, k_ON_ =1, k_OFF_ = 19.5, k_cat_ = 0.5, δ_P_ = 0.01, and k_in_ = 5/b (solid lines) or 0.357/b (dashed lines).

The variance in protein numbers at steady-state is, 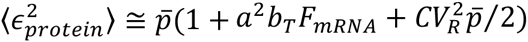, where the parameter *a*^2^ = [*K_m_*/(*K_m_* + *m*)]^2^ is the squared reaction order and, 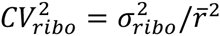 is the ribosomal noise (Sup. Mat.). The translational burst parameter, *b_T_*, is the number of proteins synthesized by a single mRNA molecule over its lifetime, which with hyperbolic translation kinetics is, *b_T_ = k_T_r*/((*K_m_ + m*)*δ_S_*). Previous work assumed first-order translation, such that *b_T_ = k_T_r*/(*K_m_ δ_s_*) (Thattai and van Oudenaarden 2001; Shahrezaei and Swain 2008). Introducing non-linearity causes the translational burst size to become a non-linear, decreasing function of mRNA abundance: all else equal, an mRNA molecule competing with fewer other mRNA’s will be translated more often in its lifetime, increasing its burst size. Alternatively, in the zeroth order regime (*m* >> *K_m_*), translational bursting goes to zero because ribosomes are so oversaturated with mRNA’s that any given mRNA is more likely to be degraded than translated. The protein noise strength and noise, then, are, respectively,

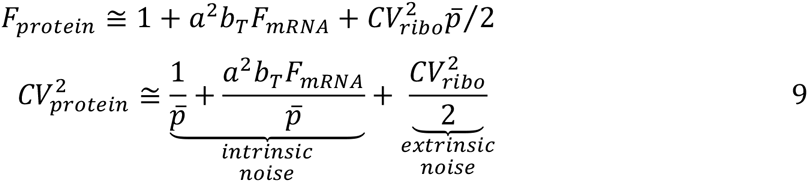

Eqn. 9 agrees well with simulation results (Figs. 2, 3) and retrieves the result of Thattai and van Oudenaarden (Thattai and van Oudenaarden 2001) (*F_protein_* ≅ 1 + *b_T_*) as a special case when there is no transcriptional bursting (*b_t_* = 1 ⇒ *F_mRNA_* = 1), no ribosome fluctuations 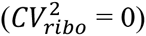, and translation is first order (i.e., linear) in mRNA abundance (*a* = 1). The first terms in *F_protein_* and 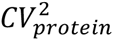 (1 and 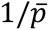, respectively) are simply Poisson terms reflecting low copy number fluctuations (Paulsson 2004; Pedraza and Paulsson 2008), while the second and third terms introduce factors that increase noise above Poisson levels. The second terms (*a*^2^*b_T_F_mRNA_* and 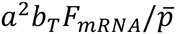, respectively) incorporate the consequences of translational and transcriptional bursting. These terms propagate mRNA fluctuations, *F_mRNA_*, to the protein level. Importantly, hyperbolic translation kinetics (0< *a* < 1) dampen the effect of mRNA fluctuations on protein noise, thus attenuating the propagation of mRNA noise via a process that might be called “hyperbolic filtering”. Together the first and second terms are considered “intrinsic noise” (Swain, Elowitz, and Siggia 2002; Raser and O’Shea 2005; Paulsson 2004; Bar-Even et al. 2006; Sanchez and Golding 2013). The last term (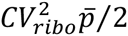 and 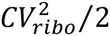) accounts for fluctuations in ribosome numbers, which create positively correlated fluctuations among protein species called “extrinsic noise” (Swain, Elowitz, and Siggia 2002; Raser and O’Shea 2005; Paulsson 2004; Bar-Even et al. 2006; Sanchez and Golding 2013). This extrinsic noise term dominates for abundant proteins (large *p*), consistent with the observation that protein noise scales inversely with protein abundance for low copy number proteins, but is independent of abundance for high copy number proteins (Blake et al. 2003; Raser and O’Shea 2004; Bar-Even et al. 2006).

Translation initiation in *E. coli* and yeast has a reaction order of *a* ~ 0.1-0.2 (3, 49), which is comfortably to the left of the protein noise peak at *a*\* = 2/3 (see Sup. Mat.; Fig 2a). This implies that, contrary to predictions based on first-order translation initiation models (Thattai and van Oudenaarden 2001; Raser and O’Shea 2005), evolving or engineering increased binding affinity of mRNA to ribosomes (i.e., reduced translational *K_m_*) will actually lower protein noise, not increase it. This occurs because reducing *K_m_* has two complementary effects on protein noise: it increases translational bursting, *b_T_*, but reduces reaction order, *a*, which attenuates noise via a phenomenon we term “zeroth-order insensitivity”. The latter effect is not accounted for in previous gene expression models, but has implications for the evolutionary and synthetic targets of noise-amelioration (see Discussion). Hyperbolic filtering also has practical implications when making inferences from data. For example, translational burst size, *b_T_*, can be estimated from Eqn 9 with knowledge of the mRNA and protein statistics; however, assuming first order reaction kinetics (*a* = 1) can lead to large underestimates of this parameter from data. For example, for *a* ~ 1/5, assuming first order translation kinetics can lead to a 25-fold underestimate of translational burst size.

### Correlation between mRNA and protein noise with hyperbolic translational filtering

We now quantify the effect of hyperbolic translation kinetics on the correlation between mRNA and protein noise. Emprical work in yeast, E. coli and human cell lines have found that there is no correlation between mRNA and protein levels in the cell, or between mRNA and protein noise (Chen et al. 2002; Ghaemmaghami et al. 2003; Taniguchi et al. 2010; Gygi et al. 1999). Here I show that this lack of correlation can be explained, in part, by modeling translation as a hyperbolic reaction.

Assuming as before that mRNA lifetimes are much shorter than protein lifetimes (δ_mRNA_ >> δ_protein_), the covariance between mRNA and protein noise, is found to be, 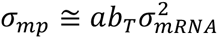. The linear regression coefficient, 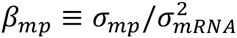 is then, *α_mp_ ≅ ab_T_*. Both of these show that hyperbolic translation kinetics (*a* < 1) diminish the statistical association between mRNA and proteins. This is most usefully illustrated by the Pearson correlation coefficient, *ρ* = *σ_mp_/σ_mRNA_σ_protein_*, which is approximately equal to,

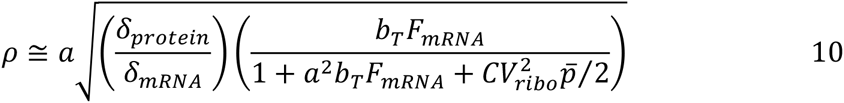

Consistent with the theoretical predictions of Tanaguchi et al (3), Eqn. 10 shows that the mRNA-protein correlation is increased with increasing transcriptional noise strength, *F_mRNA_*, is diminished with larger differences in lifetimes of mRNA and protein (*δ_protein_/δ_mRNA_* < 1), and is diminished by extrinsic noise 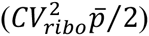. In addition, the covariance, linear regression coefficient, and correlation between mRNA and protein fluctuations all increase with increasing translational bursting, *b_T_*. The most important novel result here is that the correlation between mRNA noise and protein noise is diminished by the hyperbolic filtering of translation (that is, when *a* < 1).

### Network architecture, feedback, and buffering

While noise slows a reaction for a given substrate concentration, network feedbacks cause adjustment to substrate levels that may in turn alter reaction rates. Specifically, if a reaction proceeds more slowly, then substrate levels accumulate, which then cause the reaction to speed up again. Accounting for this feedback is crucial in quantifying the costs of noise. We now ask: How exactly does this feedback influence the total flux through noisey reaction networks? To address this problem, we now analyze how network architecture creates feedbacks that alter chemical flux, potentially compensating for noise-induced slowdown. We treat three limiting cases: 1) *h* = 1, *δ_S_* = 0; 2) *h* = 1, *δ_S_* > 0; and 3) *h* = 0.

*Cases 1* (*h* = 1, *δ_S_* = 0): *Stochastic substrate amplification and buffering:* One important feedback in noisy networks is the amplification of substrate concentration by noise (Eqn. 6), which has been studied in a process called “stochastic focusing” (Paulsson, Berg, and Ehrenberg 2000). Eqn. 6 shows that stochastic amplification will occur whenever substrate is consumed by the reaction (h = 1), as in metabolic reactions. Consequently, this steady-state adjustment of substrate creates intrinsic buffering of reaction flux that ameliorates noise-induced slowdown: the reaction is slowed by noise, allowing substrate to accumulate, which then speeds up the reaction. The extent of buffering depends on the substrate dilution parameter, *δ_S_*. In the MM network studied by Grima (Grima 2010; Grima 2009b), for example, the substrate entered the reaction volume at rate *k_in_* and exited only once consumed by the focal reaction at rate *V*. Flux balance requires that *V* = *k_in_* in such a system, so that the reaction rate in steady-state is unaffected by noise, as found by Grima (Grima 2010; Grima 2009b). Thus, under the flux constraint *V* = *k_in_*, stochastic amplification completely buffers the system against noise-induced slowdown (Fig 3).

*Case 2* (*h* = 1, *δ_S_* > 0): *Incomplete amplification and buffering.* However, the precise balancing of *V* with *k_in_* does not apply in living cells, where substrate exits the system via multiple alternative channels (δ_S_ > 0), primarily dilution by cell division, but also through excretion, titration by other reactions and enzymatic degradation. Extracellular substrate also experiences non-zero *δ_S_*, as it is diluted by bulk flow (such as in a chemostat) or by uptake from neighboring cells. Accounting for these effects modifies the flux constraint to: *k_in_* = *V* + *δ_S_s.* With this network architecture, stochastic amplification compensates for some of the diminution of *V* by noise, but not all. Consequently, stochastic amplification fails to completely buffer metabolic reactions in living cells against noise-induced slowdown.

*Case 3* (*h* = 0, *δ_S_* > 0): *Gene expression is especially sensitive to noise-induced slow-down; no stochastic amplification, no buffering.* The *V* = *k_in_* flux constraint is entirely abolished in template-mediated reactions (*h* = 0), such as protein synthesis via translation of mRNA. Because mRNA is not consumed by translation, steady-state mRNA concentration is set by the flux constraint *k_in_* = *δ_S_s*, which is independent of translation rate, *V*. Thus, stochastic amplification does not occur in translation and so cannot buffer translation from mRNA noise (Fig. 3). As a consequence, gene expression is unusually sensitive to noise.

### Cell fitness

Noise slows biochemical reactions, but how does it affect fitness? We now consider a reaction network representing a coarse-grained model of cell growth. We then analyze this reaction network in the stochastic case using the method developed above in order to quantify the effects of gene expression noise on the average rate of cell division.

During balanced exponential growth, it has been shown empirically that fitness, here defined as the division rate of a cell, is equivalent to the total rate of protein synthesis by translation in a cell (Shahrezaei and Marguerat 2015; Scott et al. 2010; Scott et al. 2014). We denote the rate of cell division by the parameter *λ*. In general, biomass synthesis involves the import of extracellular nutrients followed, ultimately, by their conversion into new biomass via translation. Translation initiation involves the binding of mRNA, in abundance *m*, to a ribosome, in abundance *r*, followed by elongation via the polymerization addition of amino acids (aa) to a growing protein chain. Elongation rate is limited by the intracellular nutrient, in abundance *s_I_*, in scarcest supply (i.e., von Leibig’s Law of the Minimum (Droop 1974; De Baar 1994; Tilman 1982)), which is typically either ATP or aa-charged tRNA’s. In eutrophic conditions, where nutrients are saturating, translation is initiation-limited (Shah et al. 2013), whereas in oligotrophic conditions, where nutrients are scarce, translation is elongation-limited (Tuller et al. 2010).

This coarse-grained cell model follows a QSSA system of ODE’s (S22) with growth rate,

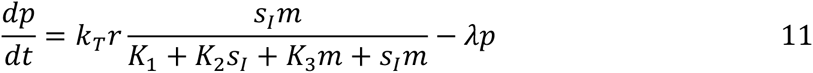

which is equivalent to a compulsory-order two-substrate enzymatic reaction with independent (i.e., non-cooperative) binding events (Ingalls 2013). *k_T_* is the translation elongation rate per mRNA per ribosome (which absorbs the number of *S_I_* per protein), *K*_1_/ *K*_2_/ *K*_3_ are affinity constants, and λ is the protein dilution rate, which during steady-state growth is equivalent to the rate of cell division. Eutrophic and oligotrophic regimes, corresponding to initiation- and elongation-limited translation, respectively, are found from this equation in the limit as *s_I_* or *m*, respectively, goes to infinity. Eqn. 11 can be rearranged to form a Monod equation, which maps growth rate to extracellular nutrient supply with a number of practical and conceptual benefits (S24).

The expected steady-state rate of cell division (cells/time) is found by setting Eqn. 11 to zero and solving for *λ*,

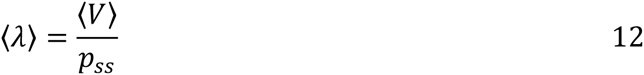

*p_ss_* is defined as the species-specific total protein content of a cell (protein/cell), which is assumed constant for a given species, *V* is the synthesis rate of proteins (protein/time).

### Fitness with noise

Following the method above, we write down a QSSA reduced system of equations for growth rate according to our coarse-grained cell model (S22), giving a system in five equations describing the time evolution of mRNA, uptake proteins, extracellular substrate, intracellular substrate and protein biomass. We assume that mRNA and extracellular nutrients undergo a linear birth-death process, supplied with burst inputs of size *b_t_* and *b_sE_* respectively, and ignore ribosome fluctuations. Intracellular nutrients are consumed by protein synthesis, but mRNA is not. Expected fitness, then, is (Sup. Mat.):

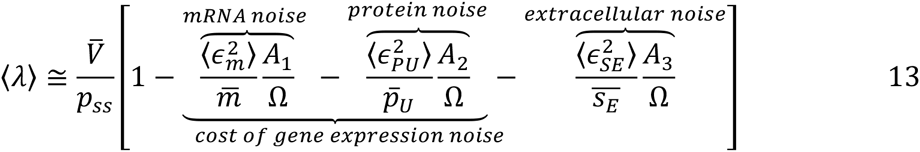

Where the *A_i_*’s are positive constants given in the SI. In reality, the protein noise term will contain contributions from all proteins involved in the pathway importing and processing nutrients to their final form utilized in translation. The MCA model of Wang and Zhang (Wang and Zhang 2011) then fits within this term. The extracellular noise term quantifies the strength of selection favoring phenotypic “bet hedging” (Starrfelt and Kokko 2012) strategies in response to environmental unpredictability.

Surprisingly, transcriptional noise directly reduces fitness. This occurs because fluctuations in mRNA levels reduce the average translation rate. This is counter to the typical view, which holds that transcriptional noise impinges on cell function only to the extent that it propagates to generate protein noise. Unlike the cost of protein or extracellular nutrient noise, which require a number of parameter estimates that are highly context-dependent, the cost of mRNA noise is easily estimated from available data if we assume that cells are growing in their eutrophic regime (*s_I_* ➜ infinity), where growth rate is limited by translation initiation. Taking the eutrophic limit of Eqn 11 gives, *dp/dt* ≅ *k_τ_rm*/(*K_m_ + m*) – *λ_p_*, which is simply a MM reaction with mRNA serving as the substrate and ribosomes as the catalyst. Noting that fluctuations in total ribosome number and mRNA number are uncorrelated, we find from equations 4, 7 and 9,

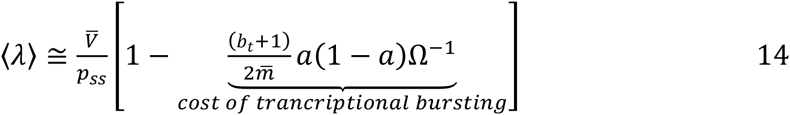

Where 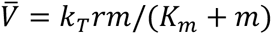. The cost of transcriptional bursting vanishes with zeroth- or first-order translation (*a* = 0 and 1, respectively) and/or as cell volume and/or mRNA abundance become large.

During balanced exponential growth of *E. coli* in rich medium: *m* ~ 1400 molecules, *r* ~ 21,000 molecules, and *k_T_* ~ 21*aa/s* ~ 4.6 proteins/min (given a mean protein length of 275*aa*). The total protein count of a cell is *p_ss_* = 2.4x10^6^ molecules. During growth in nutrient rich medium, (1 - *a*) ~ 0.85 (Arava et al. 2003; Ingolia 2014; Zenklusen, Larson, and Singer 2008). Applying these parameter estimates to 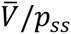 gives a doubling time for *E. coli* of ~30 minutes. Similar calculations with parameters for the budding yeast *Saccharomyces cerevisiae* (*m* ~ 15,000-60,000, *r* ~ 243,000, *k_T_* ~ 15*aa/s* ~ 2 proteins/min (average protein length 450aa), *p_ss_* = 5x10^7^, *a* = .85) gives a doubling time of ~120 minutes. Both values are in close agreement with experiments in nutrient rich media, providing an important benchmark.

In an *E. coli* cell, according to Eqn. 14, a mutation causing one unit increase in the total mRNA Fano factor results in an average loss of 7.5 proteins/min, 223 proteins/cell division, and 1.5x10^−4^ cells/generation, such that selection against mRNA noise is more than 10,000 times stronger than random drift in this species (the effective population size determines the strength of random allele frequency fluctuations and is *N_e_* = 2x10^8^ in *E. coli*). In contrast, the same mutation in yeast, which has 40-fold higher *m*, has a cost of 2.1x10^−6^ cells/generation, which is only about 20 times stronger than random drift (*N_e_* = 1x10^7^ in *S. cerevisiae*). This illustrates how fitness magnifies small differences in reaction rates by integrating over the whole cell and cell lifetime, and highlights the importance of cell volume (or more directly, molecule number) in influencing the efficacy of selection against noise.

### Rapid decay of selection efficacy with cell volume

As we just showed, the cost of noise depends on reaction volume. Because unicellular species differ over several orders of magnitude in their cell volumes, it may be possible to use these two facts in order to make predictions about taxonomic patterns of gene expression noise.

The fate of a mutation that modifies cellular noise depends on the relative magnitude of selection and random genetic drift. Selection efficiently overcomes random drift provided that 2*N_e_*|*s*| > 1 (Kimura 1957; Kimura 1962). The boundary at 2*N_e_*|*s*| = 1 defines the” drift barrier” separating efficient and inefficient selection regimes (Sung et al. 2012; Lynch 2007b). The drift barrier acts as a kind of evolutionary attractor preventing organismal perfection: recurrent deleterious mutations with effects smaller than 2*N_e_*|*s*| push species away from their adaptive optima, and selection is too weak to push them back. Thus, the balance between recurrent mutation, drift and selection establishes a suboptimal phenotypic steady-state. We now derive a scaling relationship between cell volume and selection efficacy demonstrating that the drift boundary for noise attenuation scales superlinearly with cell volume.

### Selection strength scales inversely with cell size

Selection strength, *s*, is defined as the difference in absolute fitness of mutant (*m*) and resident (*r*) alleles scaled by the mean fitness of the population. For a rare mutant we have: *s* = (*λ_m_* – *λ_r_*)/*λ_r_*. This is simply the difference in the absolute number of daughter cells produced by the mutant per resident doubling. For simplicity, we assume a haploid population and that the mutant and resident genotypes differ only in transcriptional noise. When the only source of noise is intrinsic mRNA fluctuations, the selection coefficient is approximately (see SI for derivation),

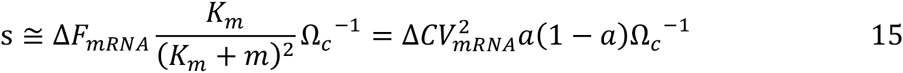

Ω_*c*_ denotes the volume of the reaction compartment. The sign of selection is determined by 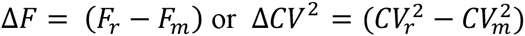, the difference in Fano factors (noise strength) or squared coefficients of variation (noise), respectively, between resident and mutant alleles. A mutation that increases transcriptional noise will be deleterious (s < 0), while a decrease in noise is advantageous (s>0). More generally, 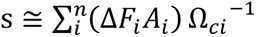, where the sum is over all *n* sources of noise affected by the mutation, the *A_i_* are constants, and *Ω_ci_* is the reaction volume of the *i^th^* noise source. Note that selection cannot act on noise levels in first or zeroth order reactions (*a* = 1 or 0, respectively), since noise does not alter the mean in these cases. Noise will, however, alter the heritability (e.g., penetrance) of a trait (Wang and Zhang 2011), which will reduce the response to selection, but we do not consider this effect here. Importantly, the strength of selection against noise explicitly depends on reaction volume.

### Drift strength scales positively with cell size

A species population size tends to scale inversely with its body size: a species must divide a limited resource pool amongst its members, resulting either in a large number of small individuals or a small number of large ones (Damuth 1981; Damuth 1987). For multicellular species the empirical scaling relationship between body mass and population size, known as “Damuth’s Law”, follows a power law with a scaling exponent of -¾, likely reflecting the ¾ power scaling of body size with energetic demands (Damuth 1981; Damuth 1987). However, in unicellular species’ metabolic rate scales either linearly (eukaryotes) or superlinearly (prokaryotes) with cell mass (DeLong et al. 2010). Assuming that *N_e_* scales linearly with N, that mass scales linearly with cell volume (i.e., cells have constant density), then *N_e_* = *B*Ω_*w*_^−*g_drift_*^, where the subscript on *Ω_w_* denotes whole cell volume and *g_drift_* = [3/4, 5/4] is the scaling exponent. The constant, *B*, is proportional to the population size of a species with unit volume, and absorbs cell density as the proportionality constant between mass and volume.

### Combining forces: Selection efficacy scales superlinearly with cell size

Combining these results implies that the efficacy of selection against noise is proportional to 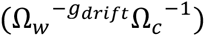. For translation, it is possible to establish a relationship between *Ω_c_* and *Ω_w_*, allowing us to reduce the number of independent variables.

Protein synthesis occurs in the cytoplasm/cytosol, suggesting that reaction-compartment volume may simply equal whole-cell volume (*Ω_c_~Ω_w_*) for translation. This relationship appears to hold between *E. coli* and yeast (*m* = 1500 and 60,000, respectively, with a 40 fold difference in cell volume) and within mammalian cells as they change volume throughout the cell cycle (Kempe et al. 2015; Padovan-Merhar et al. 2015). However, Gillooly et al (Gillooly et al. 2005) found that RNA concentration scales as body mass to the -1/4 power over a broad range of eukaryotes, corresponding to RNA number scaling with mass to the ¾ power. Although unicells were underrepresented in their sample, this agrees well with the scaling of mRNA between *E. coli* and the amoeba *Dictyostelium discoideum* where *m* = 155,000 and cell size is approximately 500-1000 fold larger than *E. coli*, such that 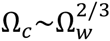 to 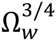.

Thus, for translation, we assume that reaction volume scales with whole cell volume according to, 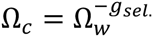, where *g_sel_* = [3/4, 1]. Therefore, the efficacy of selection against mRNA noise can be written as an allometric scaling law:

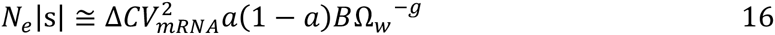

The superlinear scaling exponent, *g* = *g_sel_* + *g_drift_* = [3/2, 2¼], makes selection against noise extremely weak in large cells. For example, all else equal, selection against a mutation that increases expression noise is thirty thousand to one million times stronger in *E. coli* than in a species, such as the amoeba *Dictyostelium discoideum*, which is about 1000 times larger (Fig. 4).

**Figure 4:**
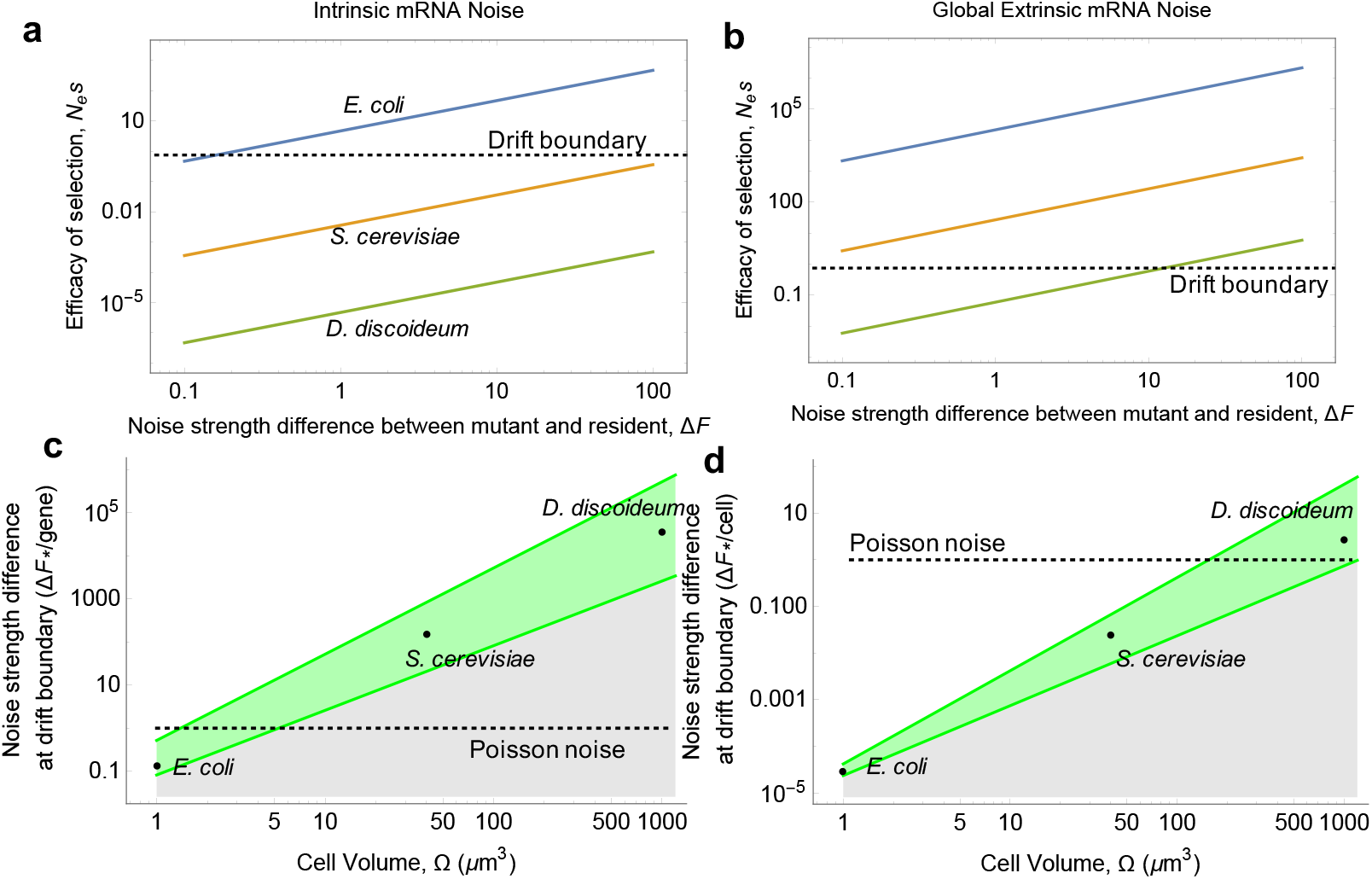
Power-law scaling of selection efficacy with cell volume. **A,B**) The predicted efficacy of selection, N_e_|s|, for *E. coli, S. cerevisae* and *Dictyostelium disoicdeum,* as a function of the jump size in noise caused by a mutation, |ΔF|, for mutations acting at a single locus (left-hand column) and mutations acting on the noise of the global transcriptome (right-hand column). The mRNA concentrations and effective population size (N_e_) estimates used for the figure are found in Table S1 (*m* = 1500, 60,000 and 155,000, respectively, and ploidy-adjusted N_e_ = 2x10^8^, 1x10^7^ and 1.1x10^5^, respectively). Reaction order was assumed to be a = 0.8 for all three species. The “drift boundary” is 2N_e_|s| =1. **C,D**) The predicted noise jump size at the drift boundary as a function of cell volume. The green region represents a range of values for the scaling exponent (-2 to - 3/2), reaction order (0.75-0.9), and in **C**) the gene number (3250-12,500). Above the green region (or individual points) selection is strong, below mutations are effectively neutral.

Solving for |Δ*F*| at the drift boundary (2*N_e_*|*s*| = 1), gives the jump size in noise strength “visible” to natural selection. If the only cost of transcriptional noise is reduced translation, then the selectively impermissible jump sizes in transcriptional noise follows the scaling relationship,

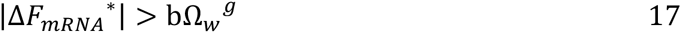

where *b* = {*K_m_* + *m*)^2^/*BK_m_*. Mutations causing jumps smaller than this critical size are “effectively neutral”, and thus invisible to selection acting on translation rate. Adaptive mutations that decrease noise by an amount less than this jump size cannot efficiently spread, while deleterious mutations with effect size below this threshold cannot be efficiently purged by selection.

Parameterizing our model with literature estimates of cell volume, effective population size, genome size, ribosomal occupancy, and translation kinetics, we predict that *E. coli* should be strongly selected to minimize intrinsic transcriptional noise simply because this noise slows translation. Whereas the cost of transcriptional noise is too weak per gene (on average) in *S. cerevisiae* or *Dictyostelium* for selection to maintain intrinsic noise at the Poisson minimum if noise-induced slowdown of translation is the only cost of mRNA noise. Fig 4 shows that in cells below about 5fL (most prokaryotes), selection favors suppression of intrinsic transcriptional noise to its theoretical minimum defined by Poisson statistics, even if the only cost of transcription noise is reduced translation rate. But because of the superlinear scaling of selection efficacy, cells only 100 times larger (most eukaryotes) cannot efficiently oppose intrinsic noise jumps even as large as Δ*F* = 1000 (again, if this is the only selection pressure for noise-minimization acting). Alternatively, jumps in global transcriptional noise (i.e., extrinsic noise) have a much greater selective effect, meaning that traits affecting extrinsic noise, such as ribosome fluctuations (Eqn 9), should be under strong selection for noise-amelioration even in moderately sized species.

## DISCUSSION

Biochemical reactions are non-linear, but are typically modeled as linear in stochastic models of gene expression due to technical limitations (van Kampen 2007). Here we have introduced a novel procedure for easily and more accurately incorporating non-linearity into models of coupled stochastic chemical reactions. We have found that hyperbolic reaction kinetics filter input noise in such a way that the average reaction rate is slowed by noise, while attenuating noise propagation. These results have a number of implications.

### A drift barrier hypothesis for gene expression noise and robustness

Cell size places severe constraints on the evolution of noise suppression. The magnitude of noise-induced slowdown of Michaelis-Menten reactions is inversely proportional to the reaction volume, which follows intuitively from the fact that, for a given molecular density, greater volumes contain more molecules and thus generate less noise. Lynch (Lynch 2007b) has previously noted the “genomic perils of evolving large body size”. Large bodied species tend to be less efficient at purging deleterious mutations due to the positive scaling of genetic drift strength with body size, leading to a number of broad taxonomic patterns including greater genomic mutation rate, transposon abundance and intron number in unicellular eukaryotes and metazoans relative to prokaryotes and viruses (Lynch 2007b; Lynch 2007a; Sung et al. 2012). Unique to gene expression noise, cell size affects not only drift but also fitness, resulting in a negative superlinear scaling of selection efficacy with cell size (Eqns. 15–17). Unicellular species span more than four orders of magnitude in cell volume (~0.1 μm^3^ Mycoplasma to more than 1000 μm^3^ in amoebazoans and other protists), with a sharp taxonomic divide separating prokaryotes and eukaryotes. “Typical” cell diameters are 0.5-5um for prokaryotes and 10-100um for unicellular eukaryotes. Taking a conservative value of a 100-fold average volume difference between prokaryotes and eukaryotes, this translates to a 1000-10,000-fold difference in the average efficacy of selection against noise for non-compartmentalized processes (Eqn. 16).

Of course, numerous other fitness effects of gene expression noise occur (Eqn. 13 and (Fraser et al. 2004; Wang and Zhang 2011; Lehner 2008; Raj and van Oudenaarden 2008; Balázsi, van Oudenaarden, and Collins 2011; Raser and O’Shea 2005)), but these must all scale with cell (or compartment) volume, such that selection for noise-amelioration will often be weaker in eukaryotes than prokaryotes. It is possible that the greater transcriptional complexity in eukaryotic cells, which employ multisubunit regulatory complexes, enhancer regions, heterochromatic gene silencing, nuclear localization into “transcription factories” and other features that greatly increase the burstiness of transcription, evolved only once greater cell size freed populations from the selective constraint of noise minimization. The cause of the size disparity between prokaryotes and eukaryotes is thought to be due to differences in cellular respiration: prokaryotes synthesize ATP via the electron transport chain embedded in the cell’s plasma membrane, which requires a high surface area to volume ratio for the whole cell in order to balance energy production and demand. Respiration via mitochondria free eukaryotes from this constraint, allowing them to achieve large cell sizes without bankrupting the cell’s energy budget. Thus, the evolution of mitochondria may have paved the way for the greater transcriptional sophistication of eukaryotes.

### Translation attenuates the propagation of transcriptional noise

Gene expression noise is generated at two levels: transcription and translation. While the molecular mechanisms controlling noise at each of these levels are becoming increasingly well understood, much remains unclear about the propagation of noise between levels. I have shown here that hyperbolic translation kinetics greatly attenuate the propagation of mRNA noise to the protein level. Given parameter estimates of translation kinetics in E. coli and yeast, the results here imply that hyberbolic translation will reduce mRNA noise by between 25 and 100 fold, substantially attenuating the propagation of mRNA noise to the protein level. The hyperbolic filtering of mRNA noise by translation also manifests as a substantial reduction in the correlation between mRNA and protein abundance (Eqn. 10). This result is consistent with data from *E. coli* showing that there is zero correlation between a gene’s mRNA noise and its protein noise (Tanaguchi et al 2010). We have shown that hyperbolic translation kinetics can help explain this lack of correlation, since the correlation coefficient between mRNA and protein noise is proportional to the reaction order parameter, *a*, which is between 0.1 and 0.2 in E. coli, implying a large reduction in the correlation. Geometrically, one can visualize this diminution of correlation by observing the Michaelis-Menten rate curve: on the flat part of the curve, all mRNA concentrations lead to the same rate of protein synthesis, thus creating a zero correlation between mRNA and protein.

A major implication of these results is that the vast majority of all intrinsic protein noise comes from translation, not transcription. Consequently, if protein noise, rather than mRNA noise, determines the functional consequences of gene expression noise, then natural selection on transcriptional noise will be extremely weak, and synthetic systems desiring control of protein noise should target translation, not transcription. We have thus come to two conclusions about transcriptional noise: 1) mRNA noise directly inhibits growth by slowing translation and 2) very little mRNA noise propagates to the protein level. These two predictions cast new light on the functional consequences of transcriptional noise in living cells.

### Survival of the flattest

We have shown that hyperbolic reactions, by filtering out high-amplitude input fluctuations, cause a reduction in both signal (average output) and noise (relative output variance) compared to first-order reactions. The strength of selection favoring organismal robustness is determined by the former, while the target of selection for increased robustness is determined by the latter. Both the strength of noise and its cost are mediated by the reaction order, *a* = *K_m_*/(*K_m_* + *s*) (Eqns. 6, 9), making this the obvious target for noise-attenuation. Noise-induced slowdown is minimal for first and zeroth order reactions (*a* =1, 0, respectively), while noise itself is minimal only in zeroth order (*a* = 0). Indeed, zeroth order kinetics completely filter out super-Poisson intrinsic input noise (Eqn. 9; Fig. 2), making the output entirely insensitive to input noise. We might call this phenomenon “zeroth order insensitivity”. Selection for robustness, then, should push reactions towards their zeroth order regime. Geometrically, the zeroth order regime corresponds to the flat part of the rate curve, such that adaptive robustness by this mechanism promotes “survival of flattest”, a term previously invoked to describe selection for robustness via very different mechanisms (Wilke et al. 2001; Codoñer et al. 2006).

There are two targets of modification for reaction order minimization: increasing substrate abundance, *s*, which moves the substrate distribution to the flatter part of the rate curve, or reducing *K_m_*, which shifts the rate curve itself. Importantly, these strategies are not equivalent (Fig. 2). Decreasing *K_m_* (while holding s constant) causes a unimodal change in product noise because *K_m_* has two complementary effects: it reduces reaction order, *a*, which filters noise, but it also increases product burst size, *b_T_* (Fig. 2). Alternatively, increasing mRNA concentration (with *K_m_* held constant) monotonically decreases protein noise because both reaction order, *a*, and translational bursting, *b_T_*, decline. This strategy increases fitness monotonically, thus avoiding the mRNA/protein noise trade-off. Importantly, increasing robustness by increasing substrate concentration is different from increasing robustness by decreasing substrate noise. To highlight this, one could force noise strength to stay constant while increasing mean substrate. The result would be increased robustness without a decrease in noise strength. This strategy undoubtedly generates other costs not accounted for in the current model, such as the energetic cost of increased mRNA synthesis. Future work should explicitly examine the evolutionary feasibility of cellular robustness via this mechanism in light of its potential trade-offs.

### Robustness via compartmentalization

The present theory provides new insight into the recent argument that compartmentalization of reactions in eukaryotic cells is a strategy for noise control (Stoeger, Battich, and Pelkmans 2016). If cell compartments are filled with reactants via active transport by membrane transporters, then the resulting hyperbolic filtering (described above) will attenuate copy number fluctuations of reactants potentially down to the Poisson minimum, though not lower (e.g., Eqns 9 and S21). But hyperbolic filtering also slows the average rate of transport, creating inefficiency. In addition, reducing the reaction volume increases the relative size of fluctuations (the noise), which slows the absolute average rate of reactions within the compartment. Future work is necessary to determine exactly when the conflicting costs and benefits of hyperbolic filtering favor reaction compartmentalization as a strategy for cellular robustness. In eukaryotes, this takes on another dimension, because mRNA is actively transported out of the nucleus and into the cytoplasm before translation. The hyperbolic filtering of transcriptional noise at this level will greatly ameliorate transcriptional noise even before mRNA is filtered by translation. Future work should model the consequences of this serial transcriptional noise filtering in eukaryotes.

### Metabolic flux

Wang and Zhang (Wang and Zhang 2011) used metabolic control analysis (MCA) to study the effect of enzyme fluctuations on metabolic flux, concluding that enzyme noise diminishes fitness in pathways with more than one enzyme. This result arises from the hyperbolic relationship between enzyme concentration and steady state flux in multi-enzyme metabolic pathways (Kacser and Burns 1973), and so, like the present paper, is a straightforward consequence of nonlinear averaging. However, MCA models assume first-order enzyme kinetics (Kacser and Burns 1973). In light of the present results, it is not immediately clear if this assumption makes the results of (Wang and Zhang 2011) an underestimate of the true cost of gene expression noise, because hyperbolic rate laws cause noise-induced slowdown at each pathway step, or an overestimate of cost, because hyperbolic rate laws attenuate noise at each step, thus damping noise propagation through the pathway. Future work should distinguish between these two possibilities.

### The crucial assumption

All of the major results of this paper rely on the validity of hyperbolic reaction kinetics as an accurate description of *in vivo* biochemical reactions. This assumption is not guaranteed, especially for translation. Translation is a unique biochemical reaction because a single substrate molecule (mRNA) is simultaneously bound to multiple enzymes (ribosomes), and it is therefore easy to imagine that translation kinetics may not be well-described by Michaelis-Menten reaction kinetics. Therefore, the present theoretical results must remain in question until future empirical work carefully measures the *in vivo* rate law of protein translation.

## METHODS

All Monte Carlo simulations were run on the elementary (microscopic) system of reactions (S17) using Gillespie’s Exact Stochastic Simulation Algorithm (SSA) (Gillespie 1977) implemented in *Mathematica* v10.3.0.0 with the open source xSSAlite package, freely available from the xCellerator project (http://www.xlr8r.info/SSA/). Each realization of the simulation is a Markov Chain with irregular (exponentially distributed) time intervals. For statistical analysis, each realization was regularized using the TimeSeriesAggregate[] command. Mean[] and Variance[] commands of the regularized time series were used to compute the moments of the Markov Chain. Data points in figures 1,2 and 3 are ensemble averages or variances over 1000-10,000 simulated realizations of the stochastic process; each realization was run for at least 10 times the half life of S or P (half-life = ln(2)/ *δ*), whichever was longer, and the value for that realization was s or *p* during the final second of the simulation run.

## Supplemental Material: “Fitness costs of noise in biochemical reaction networks and the evolutionary limits of cellular robustness”

### Derivations of Theoretical Results

#### 1.a. Stochastic Michaelis-Menten quasi steady-state approximation (MM QSSA)

Consider the canonical bimolecular reaction scheme representing the catalyzed conversion of substrate into product (3, 4):

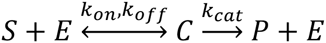

Substrate *S* reversibly binds catalyst *E* to form a reaction intermediate complex, *C*, that then decays irreversibly into *E* and product *P*. Irreversibility is allowed because living systems exist far from thermodynamic equilibrium. Because three or more body collisions are rare, all catalytic reactions can be decomposed into a series of bimolecular reactions of this type, which is why this reaction scheme provides the cornerstone of biochemistry.

This microscopic description is converted into a system of first-order non-linear differential equations (ODE’s) by applying the law of mass action. This system cannot be solved explicitly, so we apply the quasi-steady state approximation (QSSA), which assumes that the timescale of intermediate (*C*) formation is fast relative to product (*P*) formation, such that *C* equilibrates to its steady-state concentration effectively instantaneously to changes in *S* (4, 5).This separation of timescales allows us to set the concentration of *C* to it’s steady-state value, giving the steady-state rate of product formation,

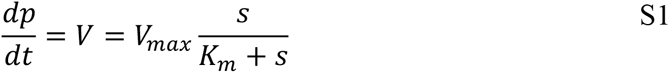

Here, *s* is reactant concentration (following convention, lowercase denotes concentration), *V_max_ = k_cat_e* is the maximum achievable reaction rate, and *k_m_* = (*k_off_ + k_cat_*)/*k_on_*, the Michaelis or half-saturation constant, is the substrate concentration at *V* = *V_max_*/2. This rate law function is a rectangular hyperbola, with *V* approaching *V_max_* asymptotically as s goes to infinite (Figure 1).

Because S1 is hyperbolic, Jensen’s inequality proves that, for random 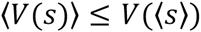. This says that the average reaction rate is less than or equal to a noise-free (i.e., deterministic) reaction with *s* = 〉*s*〈. This is shown by Taylor expanding the Michaelis Menten reaction rate about the average value of s. First, multiply the numerator and denominator of S1 by the reaction volume, Ω, to obtain a description in terms of particle numbers, *n_s_* = 〉*n_s_*〈 + *ϵ_s_*, where (*n_s_*) is the ensemble average number of species *S*, and *ϵ_s_* is a random variable with finite variance and zero mean. Expanding S1 in powers of *ϵ_s_* and taking the expected value gives,

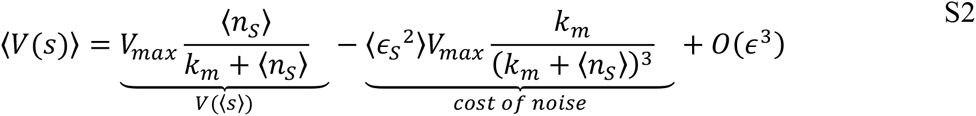

Where *k_m_* = *ΩK_m_*. The variance in *V* is,

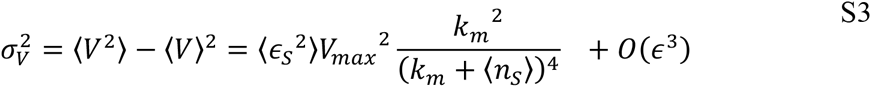

Substrate noise reduces the rate of biochemical reactions. The difference between the average stochastic rate and deterministic rate grows monotonically with increasing noise, and equality occurs only with zero noise or zero curvature of *V*(*s*).

For gamma distributed *n_S_*, the probability density function of the Michaelis-Menten reaction rate (for *V* < *V_max_*) is exactly,

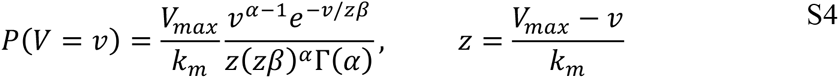

which has mean,

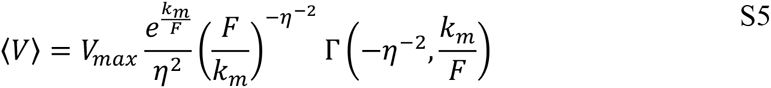

where Γ(.,.) is the incomplete gamma function. Equation S4 is verified by Monte Carlo simulations implemented with Gillespie’s Exact Stochastic Simulation Algorithm (SSA) (1) (Fig. 1).

Figure 1 shows that the effect of noise on MM-type reactions is to increase the observed Michaelis constant. Therefore, a stochastic formulation of S1 that accounts for all stochastic effects of fluctuating s at steady-state is obtained simply by substituting an “effective”, stochastic Michaelis constant using equations or S5, respectively:

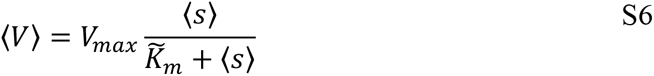

where,

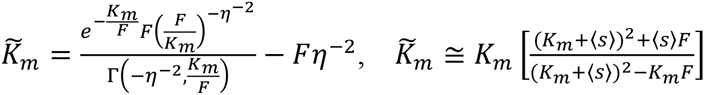

Note that 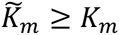 and 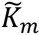 increases with increasing noise strength (measured by the Fano factor, *F*. This simple substitution retains the form of the classical MM equation (Equation S1) and provides a simple, phenomenological short-hand for the effect of noise on MM reaction kinetics.

#### 1.b. Reaction networks

##### 1.b.i The multivariate system-size expansion of the chemical master equation

We now solve for the time evolution of the moments (mean and variance) for each species in a chemical reaction network. We employ the chemical master equation and its expansion in orders of the inverse square root of system size, Ω. The chemical master equation describes the time evolution of the probability distribution of chemical species. Because these equations include bimolecular reactions they are non-linear, and are typically not explicitly solvable. The system size expansion is an approximation method, where the parameter Ω provides an objective criterion for truncation of the series expansion allowing for moment closure. For thorough reviews and extensions of the method see van Kampen (6), Elf and Erhenberg (7), and Grima (8). van Kampen introduced the system size expansion, Elf and Erhenberg generalized the system size expansion to systems of chemical reactions, and Grima extended this further by deriving a multivariate system size expansion that retained terms beyond the linear noise approximation (LNA). Here we build on this work to provide a novel method that accurately models the statistics of gene expression and metabolic networks in living cells.

We define 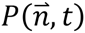 as the joint probability density function of particle numbers for all N species in a system of R reactions. Its time evolution follows the chemical master equation (7),

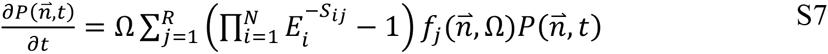

where 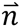 is the vector of particle numbers of all species, Ω is the reaction volume, 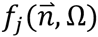 is the transition rate between states for the *j*th reaction, *S_ij_* is the stoichiometric coefficient for the net reaction, and *E* is the step operator which acts as follows: 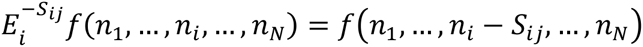. The particle number is then written as a random perturbation about the macroscopic concentration (thus translating from the extensive variable *n* to the intensive variable *x*) according to the ansatz that the relative perturbation size scales as the inverse square root of the reaction volume, in accordance with the Law of Large Numbers, leading to the substitution:

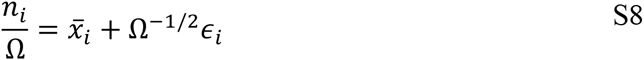

where 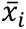 is the macroscopic steady-state concentration of species *i*, and *ϵ_t_* is a continuous random variable. Substituting into Eqn. S7 and simplifying (6, 7),

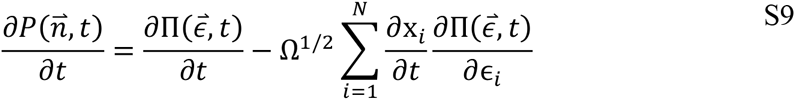

Taylor expansion of the transition rate for the *j*th reaction, 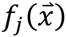, about the macroscopic rate gives,

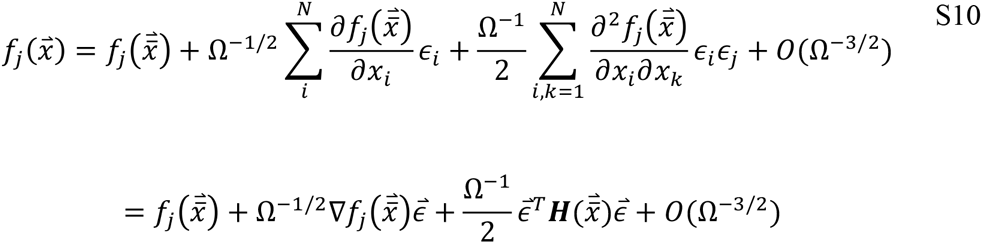

where *∇* is the gradient operator and ***H*** is the Hessian matrix.

Note that S10 differs from that of Grima (equation 13 in (8)), who included an additional term accounting for the special case of homodimerization. While Grima’s expansion works for all elementary reactions, it gives erroneous results when applied to bimolecular reactions written in QSSA reduced from. We therefore use Eqn. S10 with the caveat that it will be inaccurate for homodimerization reactions at low copy numbers. Taylor expansion of 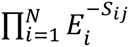 to order *O*(Ω^−3/2^) gives the differential operator (6–8),

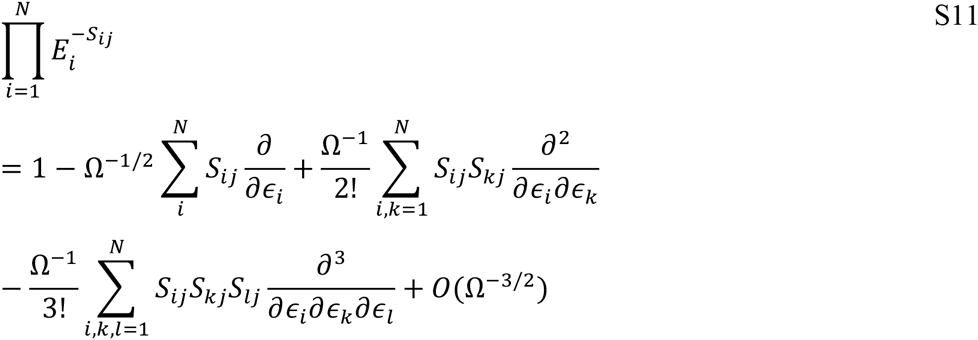

Combining S8-S11, and ignoring third derivatives (which vanish for first and second moments) we obtain,

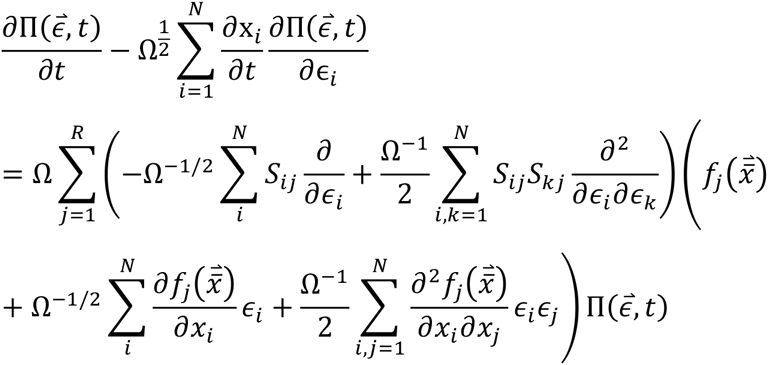

Noting that terms of order *O*(Ω^1/2^) cancel, after some algebra we arrive at the following multivariate master equation,

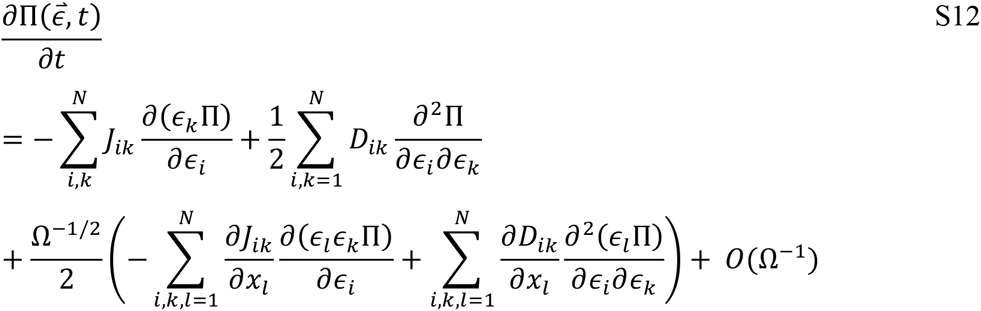

where 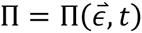, and

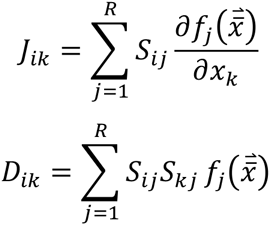

The first two terms of S12 (of order Ω^0^) together give the standard linear Fokker-Planck equation which is the sum of directional motion (e.g., convection, transport, drift) and diffusion, respectively. The terms of order Ω^−1/2^ are the nonlinear, finite volume corrections to these terms, which vanish for linear reactions and/or reactions in large volumes (i.e., in the macroscopic limit).

The stationary solution of the linear Fokker-Planck equation is a Gaussian distribution. We expect that if the relative fluctuation size is small compared to the curvature of the transition rate, then the stationary solution of S12 remains approximately Gaussian even in finite volumes with non-linear transition rates. Of course, the actual product distribution of a reaction with a hyperbolic rate law is positively skewed (and a sigmoidal rate law is leptokurtic). Assuming relatively small fluctuations, only the first two moments of 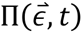 are needed to approximately characterize the full distribution in small volumes.

The time derivative of the first moment of the ith species is found by multiplying both sides of S12 by *ϵ_i_* and integrating over all *ϵ*:

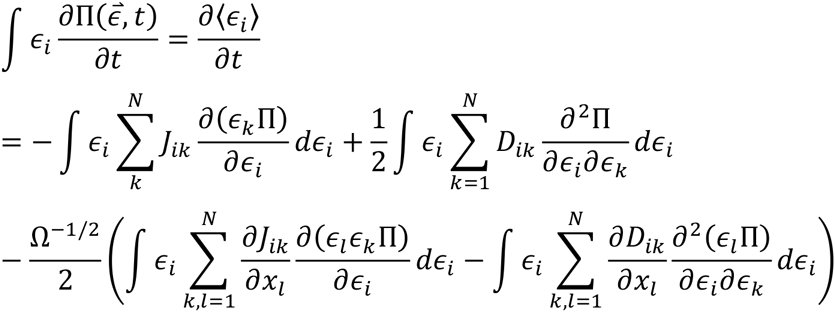

This is straightforwardly computed using integration by parts (iteratively for the second derivative terms) and applying the definitions of the mean and variance, leading to:

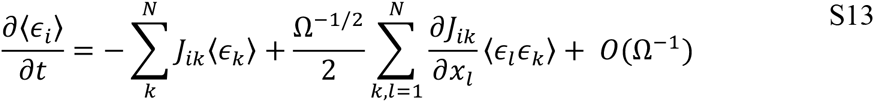

which has matrix form:

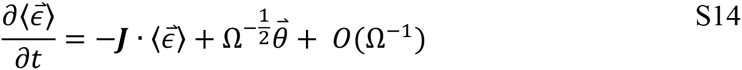

where 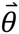 is a vector with k^th^ element:

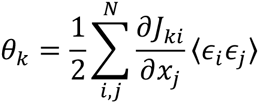

Substituting 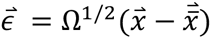, setting S14 equal to zero and solving for the average concentration vector, 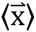, finally gives the matrix equation for first moments of the steady-state system:

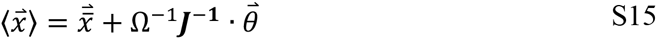

Thus, one can solve for the average of all species in a discrete, stochastic system with finite volume directly from the continuous, deterministic system of equations for the system.

One solves the second moments of the system using the same approach. One multiplies S12 by 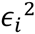 integrates (again, using integration by parts), then solves for 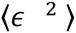 when the distribution is stationary 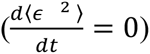. The full solution is a function of the first three moments of the joint distribution, such that the moment hierarchy does not close. Further, solving the second moments of the linear multivariate Fokker-Planck equation is already computationally intensive, without consider higher order corrections. Thus, we will ignore the non-linear finite volume correction to the second moments for the purposes of the present paper. Simulation results suggest that this leads to small error (Fig. 2, 3 main text). Therefore, we can solve for the second moments by solving the Lyapunov Equation (7):

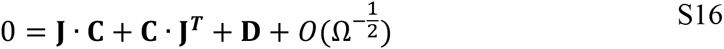

for the covariance matrix C which has elements,

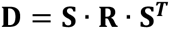

This can be solved symbolically using the *Mathematica* (Wolfram Research) command: LyapunovSolve[**J,-D**]. **D** is the noise-generating diffusion matrix given by,

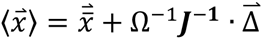

**S** is the stoichiometric matrix of the network and **R** is the matrix of the macroscopic rate function.

**Comparison to Grima (2010):**

Grima (8) previously derived results for a multivariate system size expansion to order *O*(*Ω*^1^), as we have just done. However, our final equations differ in two ways, with both quantitative and qualitative disagreement. Grima’s master equation does not include finite volume non-linear corrections to the diffusion term, which we include here only for the sake of generality and completeness, although it disappears for first moments and we neglect it for our computation of second moments. However, the following difference prevents Grima’s method from being applied accurately to a QSSA reduced system: Grima’s (8) first moments are solved using:

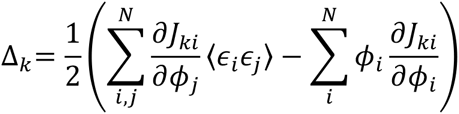

where,

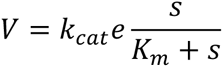

Compare this *Δ* to our *θ* above. Grima’s expression is valid when applied to all systems of elementary reactions, where the second term in *Δ_k_* accounts for homodimerization reactions (since homodimerization rates are proportional to *n*(*n*-1), not *n*^2^). However, Grima’s expression does not work when applied to reduced (e.g., “elementary complex”, pseudo-first order, biochemical) reactions. As a trade-off, the present approach is inaccurate when applied to homodimerization reactions in small volumes.

##### 1.b.ii Application to a general enzyme-catalyzed reaction in an open system

We consider the following generalized MM reaction scheme:

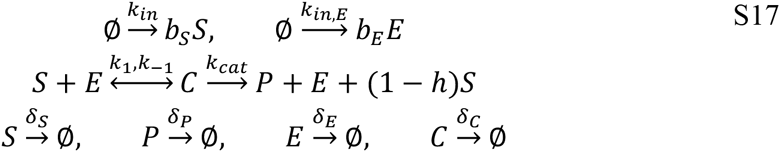

This scheme generalizes the three cases of interest. The parameter *h* is an indicator variable: *h* = 1 in reactions where S is consumed (as with most metabolic reactions) and *h* = 0 for template-mediated reactions (such as when *S* is mRNA (i.e., translation) or a gene (i.e., transcription)). When *δ_S_* = 0 the only output of *S* is via the reaction catalyzed by *E*; this corresponds to the case modeled by Grima (8) who found that noise does not alter pathway flux. This scheme corresponds to the following system of ordinary differential equations (ODE’s):

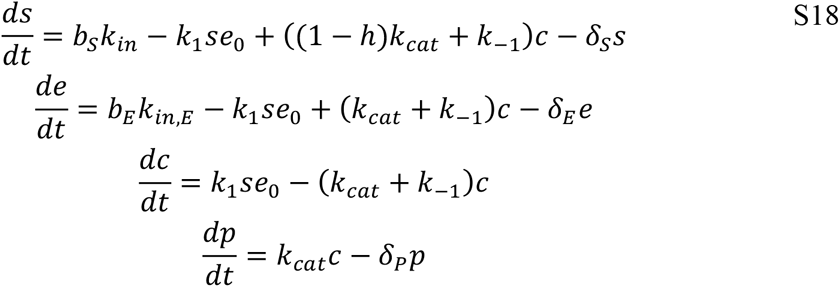

Following convention, lower case *s, e,* and *p* denote the concentration of species *S, E* and *P*, respectively. The subscript on *e*_0_ is used to make clear that this refers to “free” (unbound) enzyme concentration. Assuming that *c* equilibrates fast relative to the rest of the reaction, we can replace *c* with its steady-state solution which leads to the following QSSA reduction of the system,

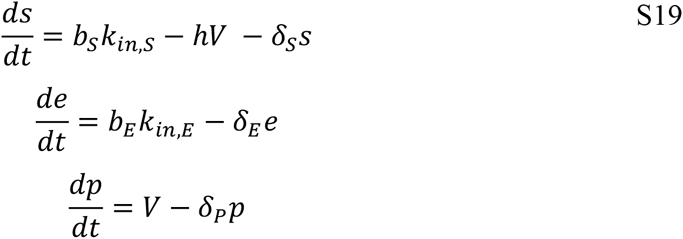

where,

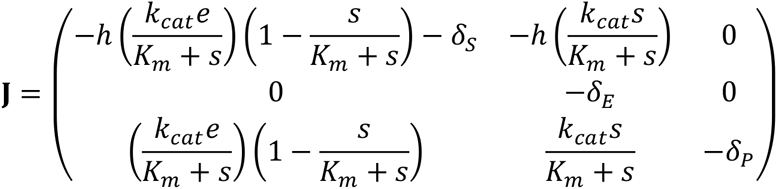

and where *e* = *e*_0_ + *c* is the total concentration of enzyme.

This system has corresponding Jacobian matrix,

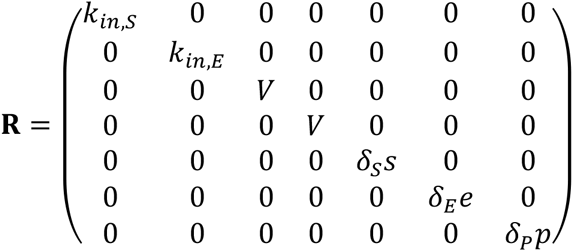

rate matrix **R**,

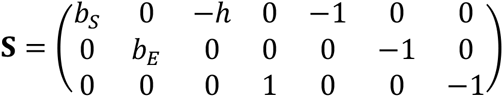

and stoichiometric matrix,

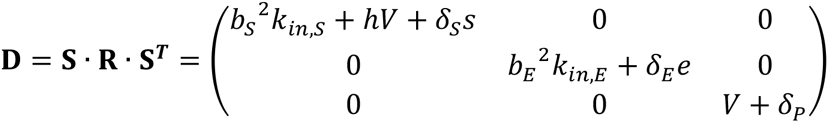

The noise-generating diffusion matrix is then,

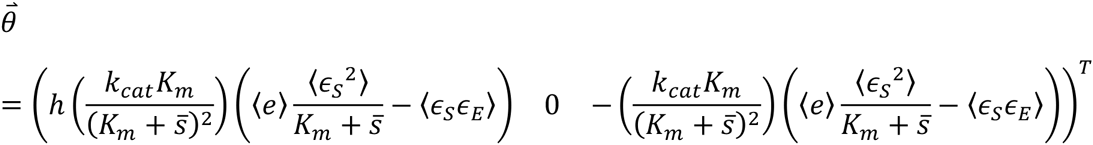

Finally, the *θ* vector equals,

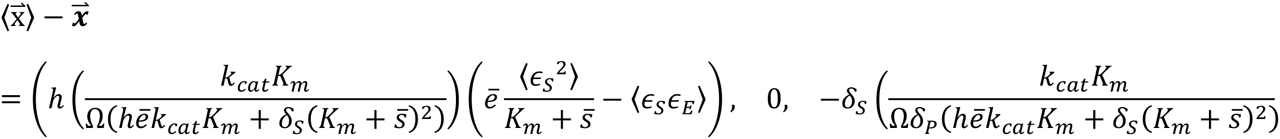

Substituting **J** and 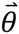 into Eqn S15 gives the noise-induced deviation of the steady-state concentrations from the macroscopic prediction:

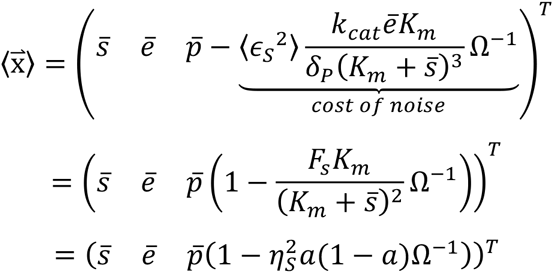

For translation (*h* = 0) the vector of steady-state concentrations simplifies to:

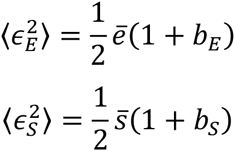

where 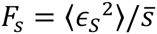 is the Fano factor of the substrate (i.e., mRNA) concentration, which equals ½(1+*b*), where *b* is the burst size of mRNA.

Four things to note from these results. First, the steady-state enzyme concentration (the second term in each vector) is not affected by noise (it has a deviation equal to 0). This is expected since d*e*/d*t* has no non-linear terms. Second, if the substrate is not consumed by the reaction (*h* = 0), then the steady-state substrate concentration is unaffected by noise (the first term in the deviation vector equals 0 when *h* = 0). Again, this is as expected since the substrate concentration when not consumed follows a simple linear birth-death process, just like the enzyme. Third, noise alters the steady-state concentration of product (i.e. the reaction flux) only if the substrate is diluted (since the third entry in the deviation vector is proportional to *δ_s_*) or if the reaction is template-mediated (*h* = 0). This is also expected: as described in the Main Text, when *δ_s_* = 0, stochastic focusing, the increase in substrate concentration due to noise, completely buffers the system from noise-induced loss of reaction flux. Finally, the cost of noise is proportional to either the Fano factor (a.k.a. the noise strength) or the squared coefficient of variation (a.k.a. the noise) of the substrate.

#### 1.c. Gene expression noise

The steady-state second moments are given by the covariance matrix, **C**. Many of these moments are too unwieldy to reproduce here. A *Mathematica* (Wolfram Research) notebook with the solution is provided as Supplementary Material. Three of the more compact moments are given here:

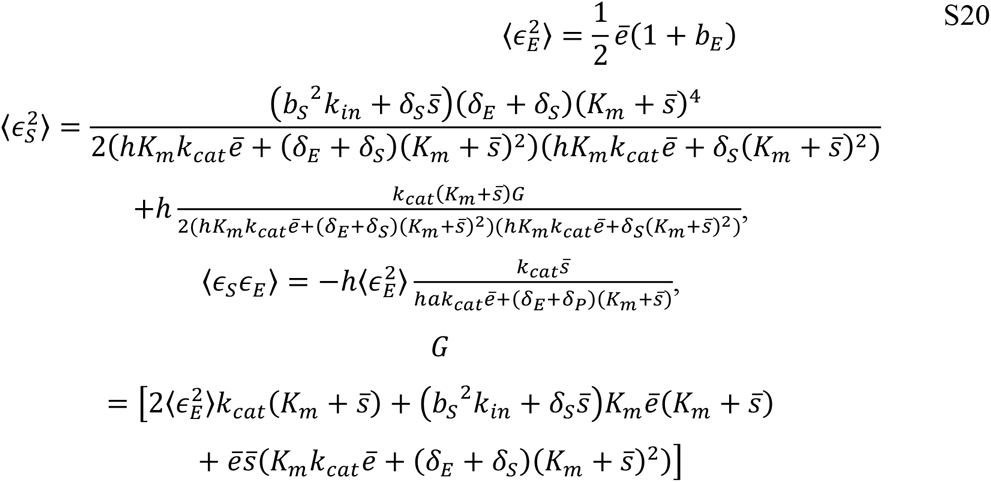

For the special case of template-mediated reactions (*h* = 0), such as translation, we find:

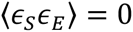

For the case when *h* =0, such as translational, the variance of product (e.g., protein) number is found to be:

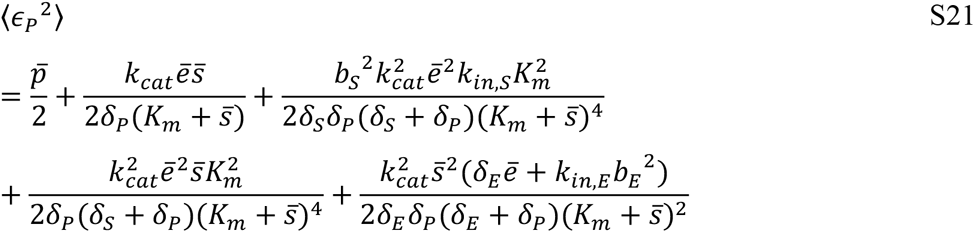

This is further simplified by noting the following:

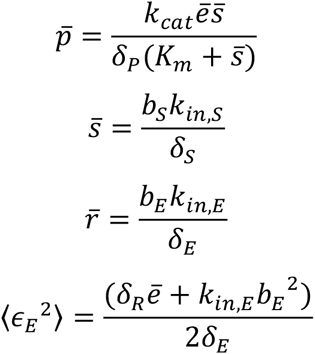

Making these substitutions and simplifying gives,

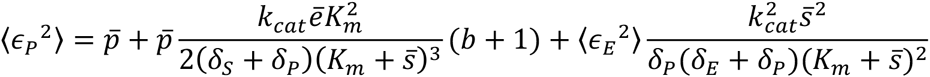

Assuming that *δ_s_* >> *δ_P_* and that *δ_E_* = *δ_P_*,

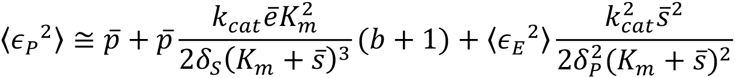

Noting that,

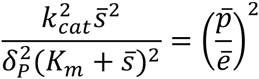

and defining the burst parameter as the number of product molecules produced in the lifetime of a single substrate molecule,

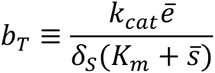

and using the definition of *a* and 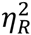 from the main text and 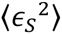 above, finally leads to,

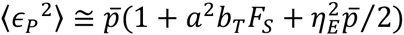

where *F_s_* is the substrate Fano factor, 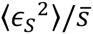.

Note that for translation with non-limiting amino acid abundance, fluctuations in ribosome concentration, 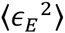, do not influence the the statistics mRNA and do not influence the mean protein abundance. Ribosome flucutations do, however, alter the variance of protein abundance, 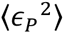. This latter effect constitutes one source of “extrinsic noise” (9, 10) in gene expression.

#### 1.d. The cost of transcriptional bursting

Substituting these statistics into the steady-state product concentration gives an expression for the effect of transcriptional bursting, *b_t_*, on protein abundance,

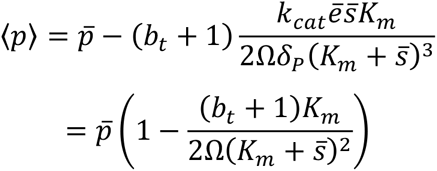

and the steady-state rate of protein synthesis 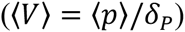

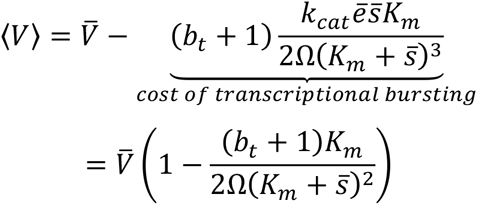

#### II.a. Response of protein noise to evolutionary changes in *K_m_* and *s*

Protein noise strength (*F_protein_*) can be adjusted in two ways: modifying *K_m_* or *s*. We now show that these strategies are not equivalent. Specifically, modifying *K_m_* results in a unimodal change in *F_protein_*, whereas modifying s has a monotonic effect. Recall from above,

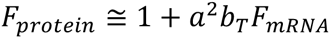

where the kinetic order parameter, *a*, is defined as (e.g., (11, 12)),

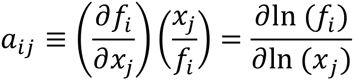

for the reaction rate *f* and concentration *x*. For Michaelis-Menten kinetics,

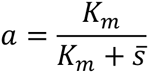

The reaction order, *a*, is equivalent to the fraction of unbound catalyst.

Evolutionary or synthetic modification of *K_m_* will alter protein noise according to,

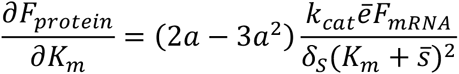

which has an interior root at,

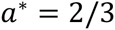

for all parameter values, which is also the maximum. Alternatively,

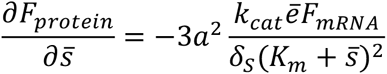

which is a monotonically decreasing function of *a*. Thus, ameliorating noise by increasing s is always a good strategy, all else equal, whereas the direction of evolution of binding affinity depends on initial conditions. Note that *a* =2/3 corresponds with weak binding affinity of an enzyme. Indeed, ribosome binding affinity ranges from *a* = 0.1-0.3, such that enhancing binding affinity will reduce protein noise.

#### II.b. Absolute fitness cost of noise

We analyze a coarse-grained cell model where cell growth relies on import of extracellular nutrient, *s_E_*, by nutrient import proteins, *p_U_*, for use in translation of mRNA, *m*, by ribosomes, *r*. Translation initiation involves the reversible binding of mRNA to ribosomes. Then elongation proceeds with the sequential addition of *n* intracellular nutrients, *s_i_*, which we define as the intracellular nutrient most limiting to translational elongation (either ATP or charged amino acids). posit that fitness, the rate of new biomass synthesis, *V*, obeys the following QSSA reduced system of biochemical equations:

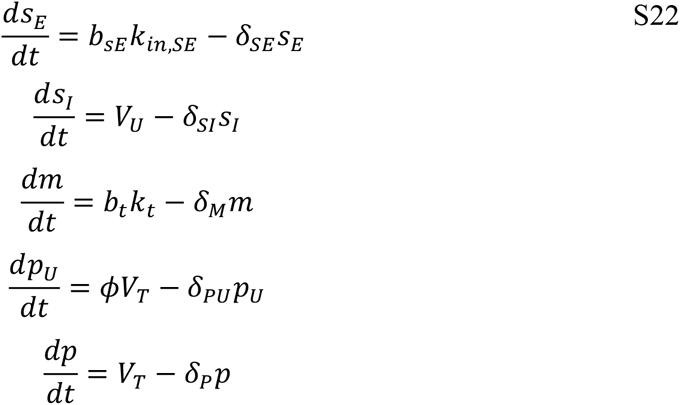

where,

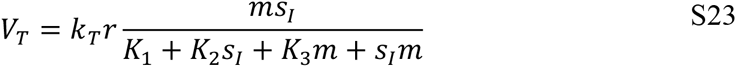

*V_T_* is the rate law for a compulsory-order (mRNA first, then *s_I_* second), two substrate (mRNA and *s_I_*) enzymatic reaction, where ribosomes are the enzyme. The parameter *φ* is the fraction of total mRNA, *m*, consisting of nutrient importer protein *p_U_* transcripts. In practice, the extracellular nutrient supply, *s_E_*, is much easier to measure and experimentally manipulate than the intracellular nutrient supply, *s_I_*. It is therefore desirable to rewrite Eqn. S23 as a Monod Growth Equation mapping extracellular nutrient supply onto growth rate (13–17). Assuming that nutrient uptake obeys MM kinetics, such that in steady-state, *s_I_* = *k_U_p_U_s_E_*/(*K_S_ + s_E_*)*δ_S_*, where *k_U_* is the catalytic constant for uptake, *K_S_* is the uptake half-saturation constant, *p_U_* is the concentration of uptake proteins, and *δ_SI_* is the dilution rate of *S_I_*. We further assume the simplest case where extracellular nutrient is used directly for biosynthesis without modification. Substituting into Eqn. S23 and rearranging gives the following Monod Growth equation:

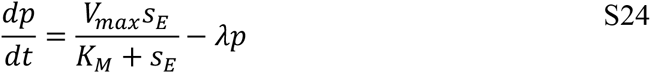

where,

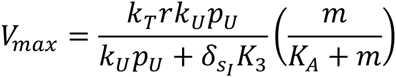

and where 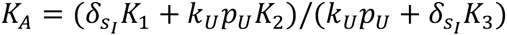 and 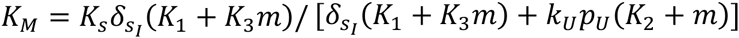. The practical and conceptual utility of the Monod equation is that it neatly disentangles cellular and environmental determinants of growth: *V_max_* and *K_M_* are intrinsic to the cell, under genetic control, and evolvable, whereas *s_E_* is extrinsic to the cell, not under genetic control and not evolvable. The version derived here unpacks the two fitting parameters, *V_max_* and *K*_M_, into their mechanistic components, revealing them both to be jointly determined by translation and uptake kinetics. Faster uptake (higher *k_U_p_U_*) increases the maximum growth rate (*V_max_*) and lowers the half-saturation constants, *K_M_* and *K_A_*, making growth faster at both high and low nutrient densities. Interestingly, this predicts an uptake-mediated negative correlation between *V_max_* and *K_M_*, which is contrary to the common view that *V_max_* and *K_M_* are positively correlated and thus in a trade-off conflict.

#### II.c. Selection strength

Absolute fitness has the following generic form if we ignore extrinsic noise:

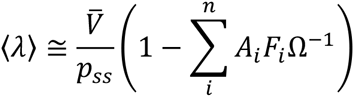

where *F_i_* is the Fano factor for substrate *i, A_i_* is a constant, and *n* is the number of substrates affecting fitness. The selection gradient on a mutation that alters fitness is,

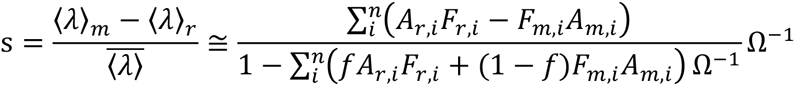

where *f* is the population relative frequency of resident alleles. Because *A* is typically small (in the standard MM reaction 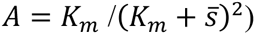, the denominator approximately equals 1 provided that *F* is not too large. For example, for *K_m_* = 350 and s = 1400 (values corresponding to translation of mRNA’s in *E. coli*), this assumption leads to only a 1% error for *F* as large as 100 and a 10% error for *F* = 1000. These *F* values are very high, and beyond any value considered in the paper. In addition, as *Ω* increases this error becomes even smaller. We therefore take this assumption as quite reasonable for the situations considered in our paper. If we further assume that genotypes differ only in noise (i.e., that there is no pleiotropic effect of the mutation), and define Δ*F_i_* as the difference in Fano factors between resident (r) and mutant (m) genotypes, then we have,

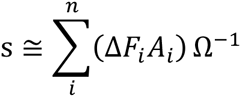

## Supplementary Figures

**Figure S1.**
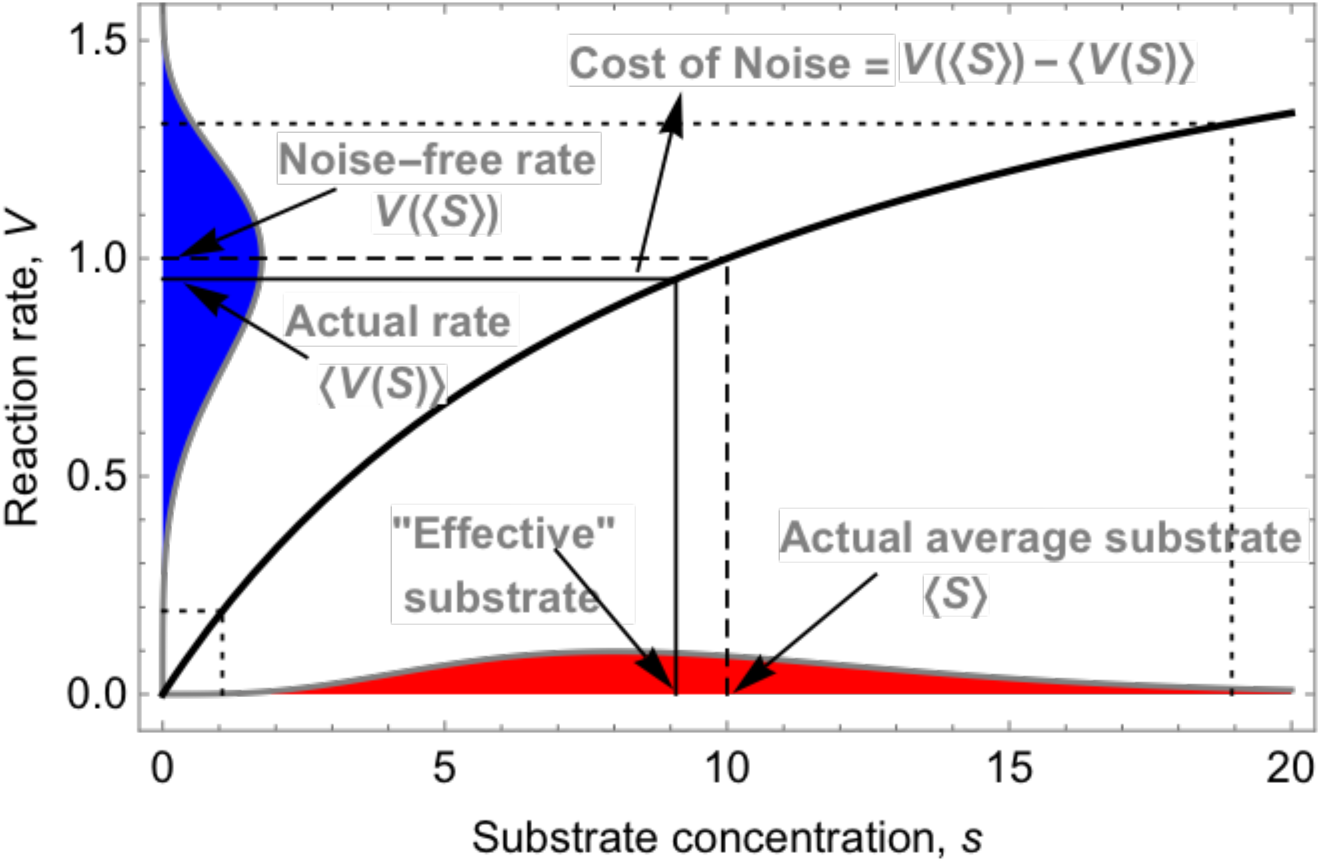
Schematic demonstrating the geometry of noise-induced slowdown for hyperbolic reaction.

**Figure S2.**
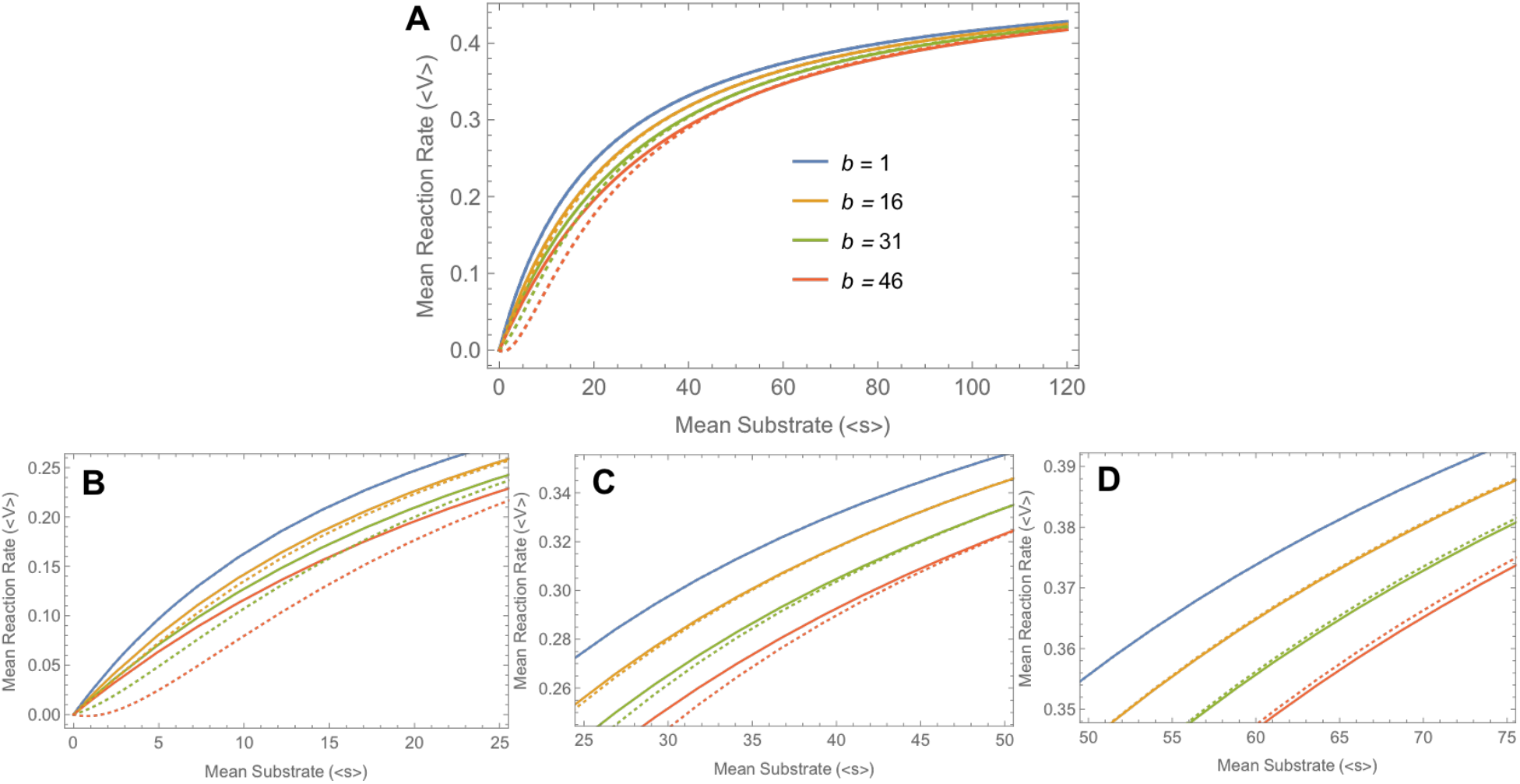
Comparison of exact and approximate predictions for noise-induced slowdown.

